# Single-cell growth rate fluctuations contribute independently from protein production to gene expression noise and decrease with average growth rate

**DOI:** 10.64898/2026.01.02.696648

**Authors:** Bjoern Kscheschinski, Dany Chauvin, Thomas Julou, Erik van Nimwegen

**Affiliations:** Biozentrum, University of Basel and Swiss Institute of Bioinformatics, Basel, Switzerland

## Abstract

Phenotypic heterogeneity is a universal feature in microbial life and has profound consequences for the behavior of single cells. However, dissecting and quantifying the sources of phenotypic noise is challenging as it requires precise quantification of the dynamics of gene expression fluctuations *in vivo*. Here, we analyze time-lapse microscopy data of single *E. coli* cells carrying fluorescent reporters for constitutive and ribosomal promoters to quantify how fluctuations in protein production, growth rate, and random sampling at division contribute to noise in protein concentrations. We find that growth rate fluctuations are largely uncorrelated with protein production and contribute substantially to noise in protein concentrations. In addition, as nutrients are varied, growth rate fluctuations decrease both in amplitude and duration with the mean growth rate. In contrast, fluctuations in production do not show a systematic dependence on mean growth rate, but exhibit surprisingly variable behaviors, even across different constitutive promoters. Moreover, although production and growth rate fluctuations are only weakly correlated, they exhibit complex cross-correlation patterns with growth rate that are both condition-and promoter-dependent. These observations suggest that, while growth rate fluctuations might be dominated by a single underlying mechanism, there are multiple sources of protein production fluctuations that vary in strength across conditions and affect different promoters to different extents. Finally, the sampling noise of proteins at cell division only contributes substantially to protein concentration noise for low-expressed genes with ten or fewer molecules per cell.

## 1 Introduction

Almost all living systems exhibit ‘intelligence’ in that they sense and respond to their environment. In relatively simple single-celled organisms like bacteria, this intelligence consists of their ability to control the expression levels of their genes and change these when environmental conditions change. However, since key molecules such as genes and mRNAs are present in very low copy numbers, the accuracy of this control is limited by fundamental thermodynamic noise at the molecular level. Indeed, it is well established that genetically identical cells grown in homogeneous environments show substantial variation in protein concentrations^1–4^. Moreover, this phenotypic heterogeneity can lead to stark differences in the behavior and responses of single cell to environmental challenges^5–10^. While limited accuracy in the control of gene expression levels may be detrimental in some situations and organisms employ strategies such as negative feedback loops to reduce noise^11–14^, in unpredictably changing environments phenotypic heterogeneity can also be beneficial for a population of microbes^15–18^.

Although the existence of gene expression noise is well established, we still understand relatively little about the mechanisms that drive gene expression noise, and in particular about the relative contributions of different noise sources to the overall noise levels of different genes. Before introducing our investigations into these questions, we place our study into context by discussing what is currently known about the sources of gene expression noise and their characteristics.

First, at the most basic level, there is thermal noise at the molecular level. The molecular components of the cell undergo erratic Brownian motion so that the precise waiting times for individual molecules to bump into each other are intrinsically random. Since genes are generally present in only one or two copies per bacterial cell, the transcription of a given gene is essentially a single-molecule event so that, even if the concentrations of all cellular components were constant, the number of transcription events per unit time would be inherently random. Similarly, for most genes, the average mRNA copy number per cell is small^4^ so that the rate of protein production, i.e., the number of translation events per unit time, is also subject to substantial sampling noise. Consequently, even two identical copies of the same gene will be expressed at different levels in a single cell due to this ‘intrinsic’ noise^1^.

It is well understood that the expression variance *σ*^2^ due to this intrinsic noise scales proportionally to the mean expression *µ_i_*^19^, i.e. *σ_i_*^2^ ∝ *µ*. It is most common to quantify gene expression noise by the coefficient of variation *σ/µ* and the contribution of this intrinsic noise to the coefficient of variation thus decreases as 1*/*√µ as the mean expression *µ* increases. However, apart from the intrinsic noise due to inherent thermal fluctuations, experimental observations show that there are additional ‘extrinsic’ sources of noise whose variance *σ_e_*^2^ scales with the square of the mean, i.e. *σ_e_*^2^ ∝ *µ*^2^. For genes with sufficiently high mean expression, their expression noise will thus be dominated by extrinsic noise. In particular, for *E. coli* it has been estimated that the expression noise of genes whose average expression is larger than 10 proteins per cell is dominated by extrinsic noise^4^.

This raises the question as to what noise sources are responsible for the extrinsic noise and why it is that the amount of extrinsic noise varies substantially from gene to gene^4,20^. In recent years, it has become clear that gene regulation itself is a major source of extrinsic noise in gene expression. Transcription factors stochastically bind and unbind promoter regions^19,21^, and fluctuations in the expression and activity of transcription factors propagate to their target promoters^18,22^. Consequently, noise levels are highly condition-dependent, more regulated genes tend to be noisier, and constitutively expressed genes are generally the least noisy, independent of growth condition^22^. In addition, as nutrients are varied, noise levels have been observed to systematically decrease with the mean growth rate of the cells, and even the ‘noise floor’, i.e., the noise level of unregulated genes with the least noise, decreases with growth rate^22^. Although the origin of this growth-rate dependence of expression noise is not yet understood, theoretical studies suggest that it can dramatically increase the benefits of gene expression noise^16^.

Here we set out to understand what mechanisms drive the extrinsic noise levels of constitutively expressed genes in bacteria, and how these vary across growth conditions. Generally, the concentration of a protein in the cell depends on its rate of production, its partitioning at the last cell division, and the global rate of dilution of proteins due to the growth of the cell. Fluctuations in any of these processes can thus drive fluctuations in protein concentration. However, to quantify the contributions of these processes requires tracking them in time. Since experimental techniques such as RNA sequencing and flow cytometry only provide snapshots of the data, they are blind to the dynamics of the underlying stochastic processes that led to the observed noise.

Therefore, to study the contributions of fluctuations in production, dilution, and random partition at cell division, we here used an integrated setup combining microfluidics, fluorescence time-lapse microscopy, and automated image analysis. In particular, we use a microfluidic device called the Dual Input Mother Machine to track single *E. coli* cells that are trapped in narrow growth channels connected to a large channel, which provides the cells with a continuous flow of nutrients and flushes away cells that are pushed out of the growth channels^23,24^. Using time-lapse fluorescence microscopy, we track growth and the production of GFP from a set of chromosomally integrated transcriptional reporters. In particular, to track the behavior of constitutively expressed genes, we chose a set of 4 synthetic promoters that were obtained by screening a library of random DNA fragments for expression activity^18^, with two promoters expressing at a ‘medium’ level (i.e. around the median of all native *E. coli* promoters, and two ‘high’ expressing promoters (around the 95 percentile of all native promoters). In addition, to contrast the behavior of these constitutive promoters with those of highly regulated genes, we also performed the same measurements with a transcriptional reporter from a ribosomal protein gene promoter and a ribosomal RNA promoter.

We obtained time-lapse microscopy data with these 6 promoters across 4 different growth media with different average growth rates, i.e. M9 minimal media supplemented with acetate, glycerol, glucose, or glucose and amino acids. To accurately infer the single-cell dynamics of protein production, growth rate, and random partitioning of proteins at cell division, we used a recently developed Bayesian inference method, RealTrace^25^, that rigorously corrects for measurement noise using maximum entropy process priors.

It is well appreciated that cell metabolism depends on the fluctuating concentrations of gene products and metabolites, which leads to considerable growth rate fluctuations in single cells^26,27^. While the impact of fluctuating abundance of molecules required for metabolism has been studied experimentally^26^ and theoretically^28^, less attention has been devoted to the question of how growth rate fluctuations affect protein concentrations^29^.

Here, we find that growth rate fluctuations are a major contributor to protein concentration fluctuations, on par in importance with fluctuations in protein production. In particular, because the instantaneous growth rate sets the intracellular dilution rate within the cell, and growth rates generally fluctuate independently of protein production fluctuations, these fluctuations are a major source of protein concentration fluctuations. Moreover, we find that growth rate fluctuations systematically increase in amplitude and duration as the average growth rate of the cells decreases.

In contrast to the systematic behavior of growth rate fluctuations with average growth rate, protein production fluctuations do not change in a systematic manner with growth rate. Instead, production fluctuations are highly variable across promoters, even across synthetic constitutive promoters that express at a similar level. Protein production rates also exhibit complex cross-correlation patterns with growth rate that vary across conditions and promoters. This surprising variability in the production fluctuations across synthetic constitutive promoters indicates that production fluctuations are likely driven by multiple mechanisms that vary in intensity across conditions and affect different promoters to different extents. Finally, like the contribution of intrinsic noise to concentration fluctuations, we find that random partitioning of proteins at cell division also only makes a substantial contribution to expression noise for genes with a mean expression below 10 proteins per cell.

In summary, our findings show that growth rate fluctuations are a major driver of protein concentration fluctuations and that these fluctuations systematically decrease with average growth rate. This may explain the general decrease in extrinsic gene expression noise with growth rate, and explain why slow-growing cells tend to be more phenotypically heterogeneous.

## 2 Results

### 2.1 Growth rate and volumic protein production rate fluctuate independently

Our observations track the size and total fluorescence of single cells through time. We first consider how to parametrize the dynamics of growth and fluorescent protein production. To illustrate our discussion, we use data from experiments where cells were growing in M9 minimal media supplemented with glucose and fluorescence quantified GFP expressed from a synthetic constitutive promoter that we denote by *hi1*. The strain is described in the Methods, and the promoter sequence is provided in the Supplementary Materials Sec. D. We processed the raw measurements using a recently developed Bayesian inference method, RealTrace^25^, which reports the estimated length *l*(*t*) and total fluorescence *g*(*t*) (including error-bars) for each cell at each time point *t*.

For growth, the situation is relatively straightforward. Single cells expand approximately exponentially through their cell cycles^23^, although the rate of growth fluctuates in time^26^. We define the instantaneous growth rate *λ*(*t*) as the time derivative of the log length, i.e.,

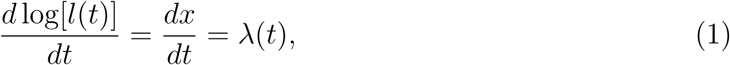

where we have defined *x*(*t*) = log[*l*(*t*)]. Note that, within a single condition, the width of the cells is roughly constant, so that cell length and volume are proportional. In the following, we will assume cell volume is proportional to *l*(*t*) and refer to cell length and size interchangeably^30^.

The time evolution of total fluorescence *g*(*t*) is determined by total fluorescence production *Q*(*t*) and loss of fluorescence due to bleaching and GFP degradation at a total rate *β*

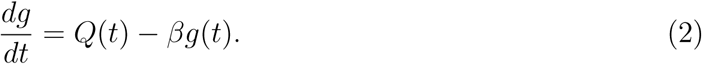

Choosing a parametrization of the production *Q*(*t*) is less straightforward. In the simplest imaginable scenario, concentrations of all relevant intracellular molecules, such as RNA polymerases, mRNAs, ribosomes, tRNAs, etc., would all remain constant as the cell expands exponentially. In this simplest scenario, the total production *Q*(*t*) would be directly proportional to cell size. A scatter of cell length against total production *Q*(*t*) indeed shows a substantial positive correlation (Fig. 1c). Moreover, defining the volumic production rate

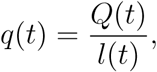

**Figure 1:**
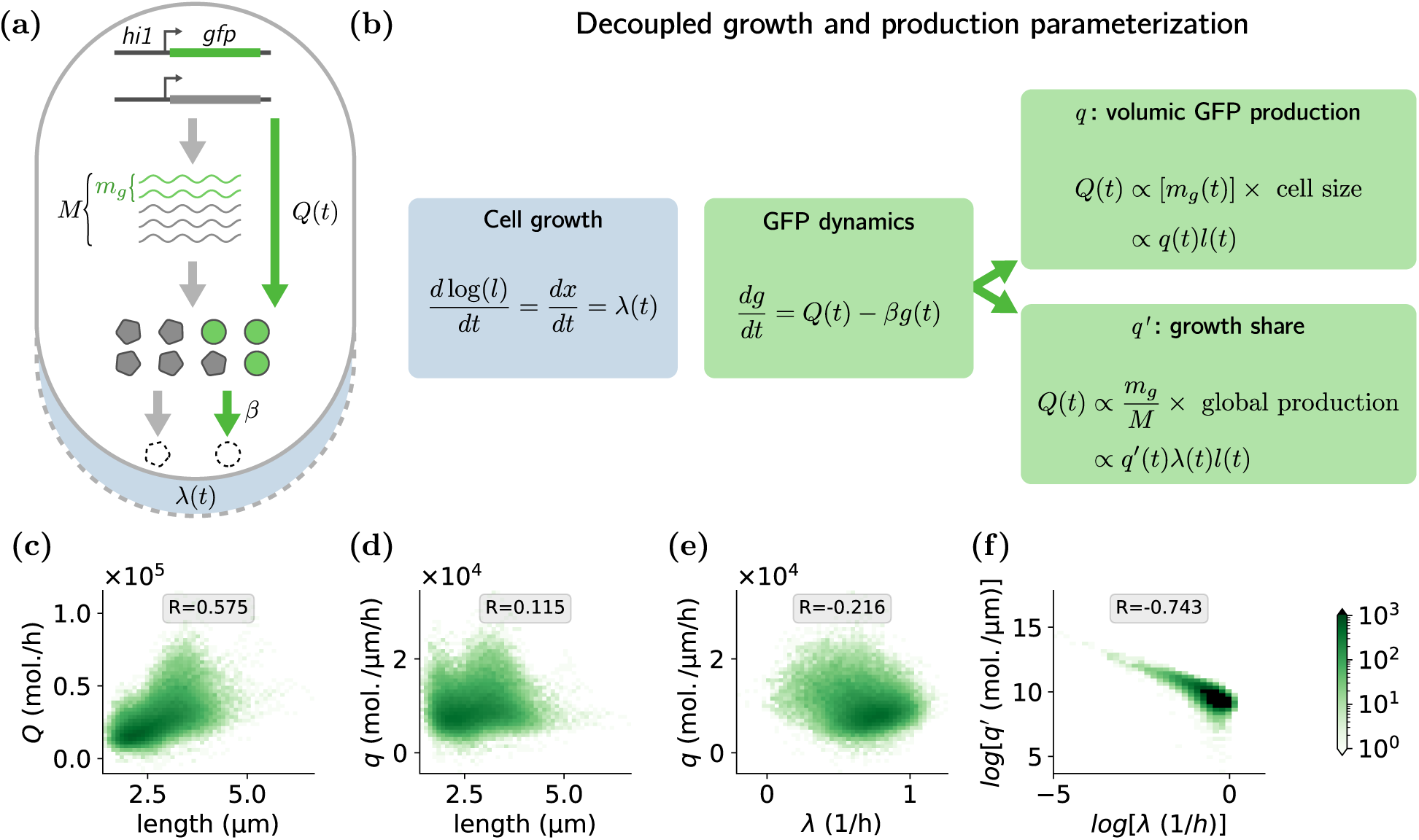
Protein production rate correlates with cell size but not with growth rate. (a) Illustration of the processes contributing to the total rate of GFP production *Q*(*t*) for a transcriptional reporter, i.e., transcription, translation, and maturation of GFP molecules. *M* corresponds to the total number mRNAs, *m_g_* the number of mRNAs for the GFP reporter, and *β* the bleaching rate. (b) The instantaneous rate of cell growth is parametrized by the time derivative *λ*(*t*) of the logarithm of cell length. The total rate of fluorescence production *Q*(*t*) can either be parametrized by production per unit volume *q*(*t*), or by production per unit of total cell growth *q*^′^(*t*). (c) The total production rate *Q*(*t*) is correlated with cell size. (d) The volumic production rate *q* has virtually no correlation with cell size. (e) A small negative correlation between volumic production *q* and growth rate is driven by a small fraction of slow-growing cells with elevated production. (f) Parameterizing the production as proportional to growth, i.e., *q*^′^ = *Q/*(*λ×l*), induces a strong anti-correlation with growth rate.

we find that there is indeed virtually no correlation between volumic production and cell size (Fig. 1d).

Besides correlating with cell size, one might also ask whether the production *q*(*t*) correlates with the rate of growth *λ*(*t*) as well. In particular, if one assumes that the total protein concentration of the cell fluctuates relatively little, then the growth rate *λ*(*t*) reflects the total volumic rate of protein production, i.e. *dP/dt* ∝ *λ*(*t*)*P* (*t*), with *P* (*t*) the total amount of protein in the cell. If, in addition, fluctuations in total protein production are mostly driven by fluctuations in the total rate of translation, then one would expect *all* genes to be translated faster, and thus the GFP production *q*(*t*) would be expected to correlate with the growth rate *λ*(*t*).

Indeed, it is often assumed that protein production is limited by ribosome availability, i.e. that the cell operates in a regime where essentially all ribosomes are translating mRNAs and mRNAs are thus ‘competing’ for ribosomes^31–33^. If we denote the translation rate by *k_t_* and the number of ribosomes by *R*, then in this regime the total rate of protein production is just *dP/dt* = *k_t_R*, and is insensitive to the total amount of mRNA *M*. Similarly, neglecting differences in translation initiation rates across mRNAs, the production of GFP is proportional to the *fraction* of mRNAs that are mRNAs for GFP, i.e.

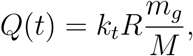

with *m_g_* the number of mRNAs for GFP. Thus, in this regime, the production of GFP *Q*(*t*) is proportional to the total protein production *k_t_R*, which in turn is proportional to both cell size *l*(*t*) and growth rate *λ*(*t*).

If cells indeed operate in this regime, we would thus expect the production rate *Q*(*t*) to not only scale with cell size *l*(*t*) but with the growth rate *λ*(*t*) as well. However, as shown in Fig. 1e, there is no positive correlation at all between volumic production *q*(*t*) and growth rate *λ*(*t*). Instead, there is a weak negative correlation resulting from a small subset of observations where cells grow very slowly and have elevated production. Indeed, if we parametrize GFP production as proportional to the total growth of cell volume, i.e.

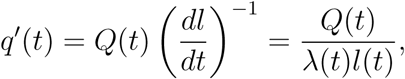

then we find that *q*^′^(*t*) is clearly negatively correlation with growth rate *λ*(*t*) (Fig. 1f). These observations show that, at least when growing in minimal media with glucose, protein production for a constitutive gene does not correlate with the growth rate of the cell, as would be expected when protein production is limited by ribosome activity.

This raises the question of what kind of regime the cell is operating in. One possibility is that the growth of cell size does not correlate with protein production, i.e., that fluctuations in growth rate *λ*(*t*) mainly reflect growth of the cell membrane and not the total protein content. Although we cannot exclude this scenario, it would imply that the total protein density in the cell would undergo substantial fluctuations. Another possibility is that the cell operates in a regime where protein production is limited not by ribosomes but by the total amount of mRNA *M*. In this regime, the protein production for each gene *g* is proportional to its mRNA copy number *m_g_* and the rate at which ribosomes initiate per mRNA. That is, if *k_i_* is the initiation rate and *R/V* is the ribosome concentration (with *V* the cell volume), protein production is given by *dP_g_/dt* = *m_g_k_i_R/V*. In this regime, the total production is proportional to total mRNA *dP/dt* = *Mk_i_R/V* and GFP production is proportional to GFP mRNA copy number *Q*(*t*) = *m_g_k_i_R/V*. If the mRNA level of GFP *m_g_* does not correlate with total mRNA *M*, then the fluorescence production *Q*(*t*) will indeed not correlate with growth rate *λ*(*t*).

In summary, based on our observations with the constitutive promoter in minimal media with glucose, we parametrize the GFP production by the volumic production rate *q*(*t*). The dynamics of the GFP concentration *c*(*t*) = *g*(*t*)*/l*(*t*) reads,

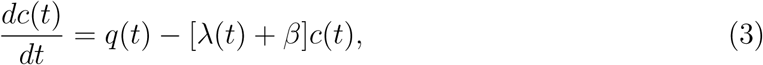

where the second term on the right-hand side corresponds to the decay of fluorescence concentration due to cell growth (at rate *λ*(*t*)) and due to both GFP degradation and bleaching (at rate *β*). Equation (3) shows that, besides fluctuations due to random partitioning of proteins at cell division, fluctuations in GFP concentration result from both fluctuations in production *q*(*t*) and fluctuations in growth rate *λ*(*t*).

### 2.2 Growth rate fluctuations are a major source of gene expression noise

While it is well appreciated that fluctuations in protein production rate *q*(*t*) contribute to fluctuations in protein concentration, it is often neglected that fluctuations in the growth rate *λ*(*t*), by causing fluctuations in dilution rate, can also contribute to protein concentration noise. To exemplify the role of growth rate fluctuations to gene expression noise, we first focus on the data discussed in the previous section, i.e., where *E. coli* cells were growing on M9 minimal media supplemented with glucose and expressing GFP under the control of the synthetic constitutive promoter *hi1*. In particular, we compared the experimentally observed dynamics with the dynamics that would result if we *only* retained the fluctuations in growth rate, i.e., if there were no fluctuations in protein production and no noise at cell division (Fig. 2).

**Figure 2:**
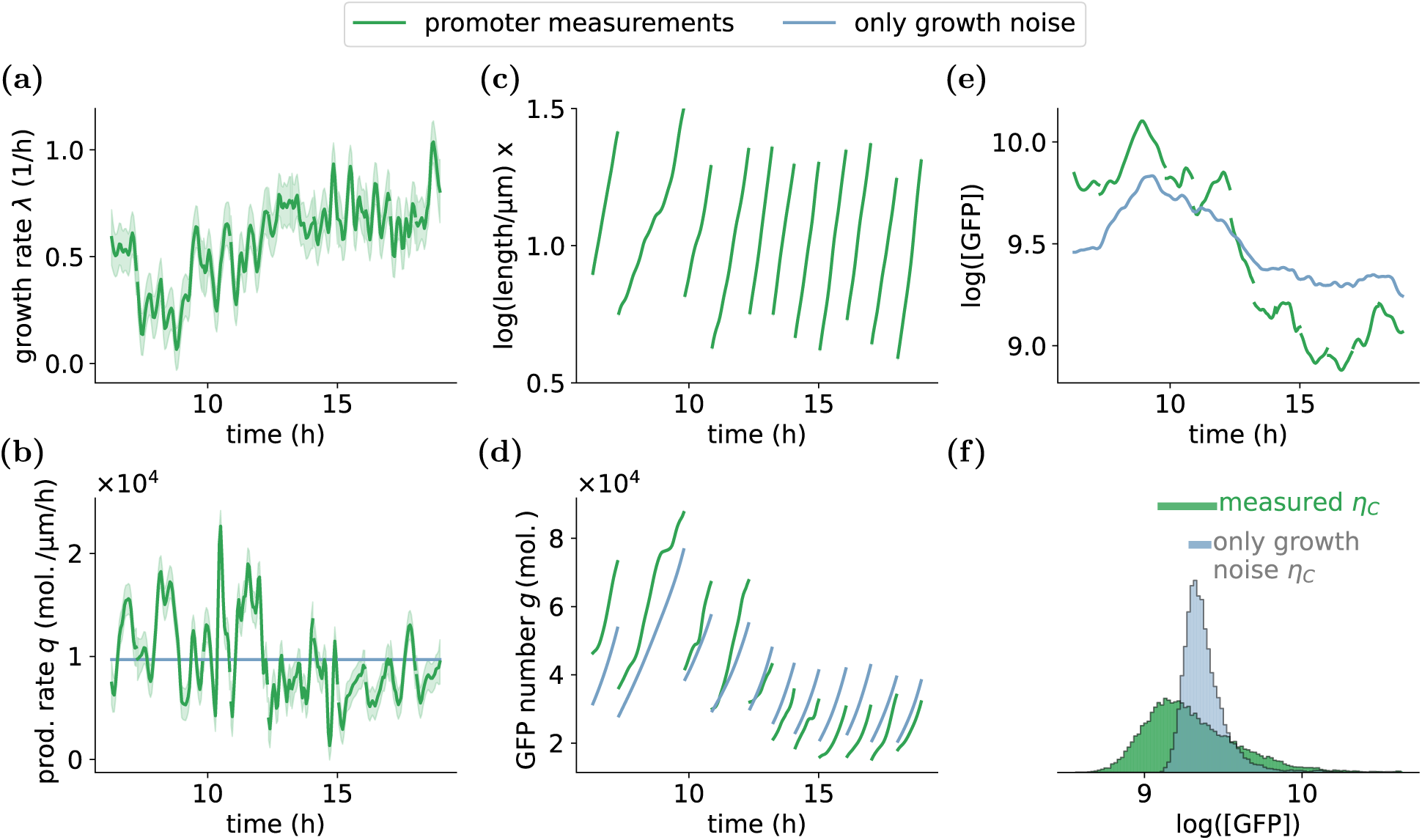
Growth rate fluctuations cause substantial noise in gene expression. Growth and gene expression dynamics for an example lineage of single *E. coli* cells growing in minimal media with glucose and expressing GFP from the constitutive synthetic promoter *hi1* (green), together with the corresponding dynamics in a hypothetical scenario in which protein production is constant, and there is no noise at cell division (blue). (a) Growth rate as a function of time. The shaded ribbon indicates the one standard deviation error bar. (b) Volumic GFP production rate *q*(*t*) as inferred from the measurements (green, with shading corresponding to error-bars) and under hypothetical constant production (blue). (c) Logarithm of cell length *x*(*t*) = log[*l*(*t*)] as a function of time. Note that there are 11 cell cycles in this example lineage. (d) Total GFP content *g*(*t*) as a function of time inferred from the measurements (green) and under constant production and absence of division noise (blue). (e) Log GFP concentration as a function of time inferred from the measurements (green) and under constant production (blue). Note that in the latter case, concentration fluctuations are solely driven by growth rate fluctuations. (f) Histogram of log GFP concentration for the entire data set (green) and the corresponding data assuming only growth rate fluctuations (blue). The bars at the top indicate the inter-quartile ranges of log GFP concentration *η_C_* for both histograms.

Our fluorescence time-lapse microscopy measurements provide estimates of the instantaneous growth rate *λ*(*t*) (Fig 2a), volumic GFP production *q*(*t*) (Fig 2b), cell size *l*(*t*) (Fig 2c) and total GFP content *g*(*t*) (Fig 2d) across time for all cells. Since variation in GFP levels generally scales with their mean, we use the logarithm of GFP concentration as a measure of gene expression (Fig. 2e), i.e., so that equal fold-changes in protein concentration correspond to equal changes in gene expression. We then compared the growth and gene expression dynamics inferred from the measurements with a hypothetical situation in which there are only growth rate fluctuations. That is, we set the volumic production rate *q*(*t*) constant and equal to the average production rate 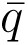 in all cells at all times, and integrate the total GFP and concentration dynamics using the cell sizes and growth rates observed in the data (blue lines in Fig 2b, d, and e). In addition, to also remove sampling noise at cell division, we divide the GFP molecules among the daughter cells exactly proportional to their sizes, so that the GFP concentration remains equal in both daughters at birth. Finally, Fig. 2f shows the distributions of log GFP concentration in the real data (green) and in the hypothetical scenario where there are only growth rate fluctuations (blue). Comparing the inter-quartile ranges of these distributions (bars in Fig. 2f), we see that the growth rate fluctuations are responsible for a substantial fraction of the total gene expression noise. Thus, at least for this high-expressing constitutive promoter in this growth condition, growth rate fluctuations are a major driver of gene expression noise. Next, we systematically investigate how the impact of growth rate fluctuations varies across growth conditions and promoters.

### 2.3 Growth rate fluctuations decrease with average growth rate

We next calculated the contributions of growth rate fluctuations to gene expression noise of constitutive promoters across four different growth conditions with different average growth rates. In particular, we tracked single-cell gene expression from two highly expressed synthetic constitutive promoters (*hi1* and *hi3*) and two medium expressed synthetic constitutive promoters (*med2* and *med3*) across four environments with different nutrient sources with increasing mean growth rates: acetate, glycerol, glucose, and glucose plus amino acids (gluc.+aa). To contrast the behavior of constitutive promoters with that of highly regulated promoters, we also tracked single-cell gene expression from one ribosomal protein promoter (*rplN*) and one ribosomal RNA promoter (*rrnB*) in the same conditions.

At the bulk population level, it has been observed that as the growth rate is varied by varying nutrients, the expression of ribosomal genes increases with growth rate, while the expression of constitutive genes decreases^34^. In line with these observations, we find that the log GFP concentrations for the four constitutive promoters all decrease as the average growth rate increases, whereas log GFP concentrations for both ribosomal promoters increase with growth rate (Fig. 3 and Fig. S1).

**Figure 3:**
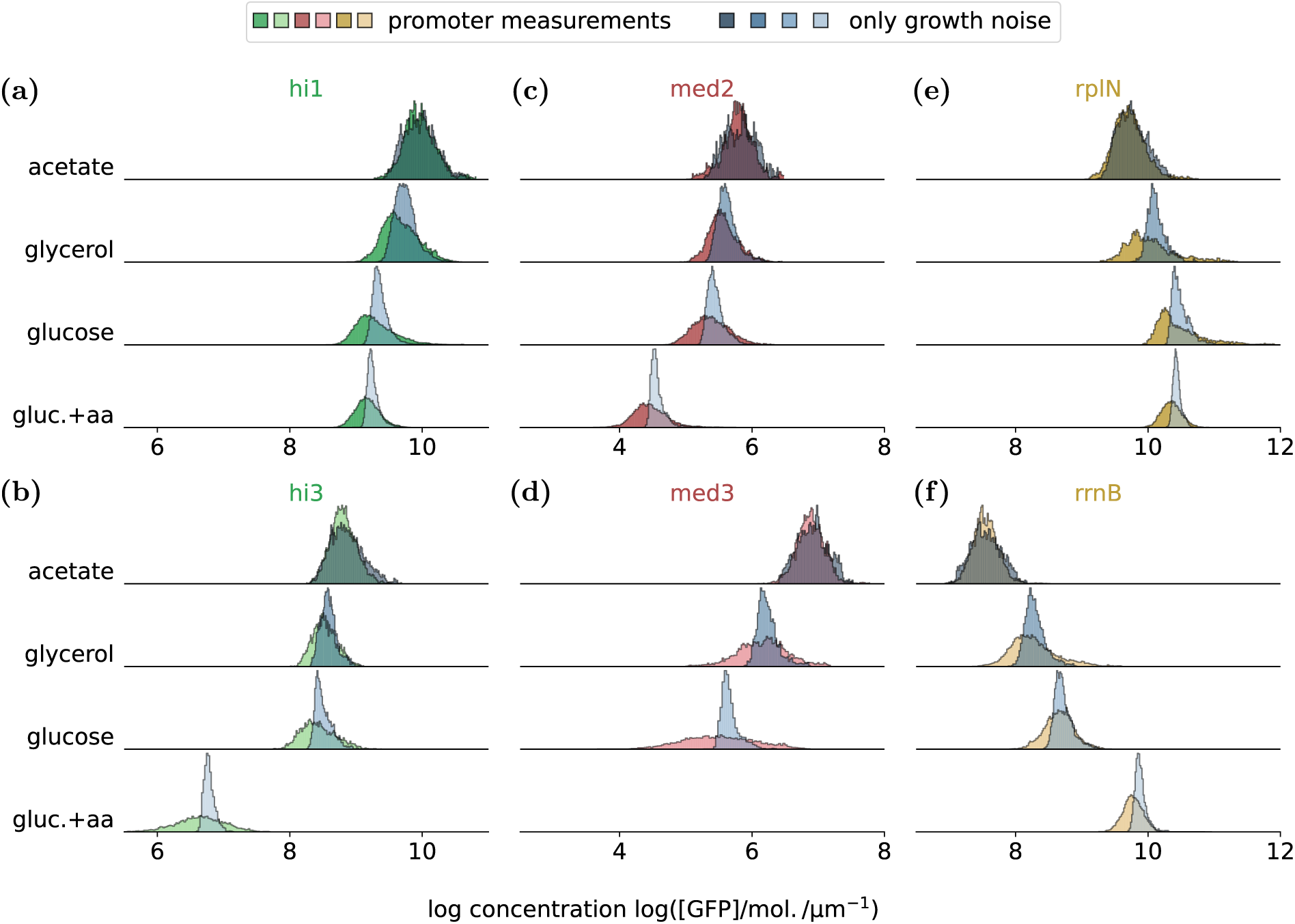
Contribution of growth rate fluctuations to gene expression noise across promoters and growth conditions. Distributions of log GFP concentration for high-expressing constitutive promoters (a-b), medium-expressing constitutive promoters (c-d), and ribosomal promoters (e-f) in four different growth conditions (rows). The blue histograms show the distributions resulting purely from the fluctuations of the growth rate, as the volumic production rate is set to a constant 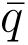 and the division noise is eliminated. The expression of the *med3* promoter in glucose plus amino acids is not shown because it is so low that, due to auto-fluorescence fluctuations, it is impossible to reliably estimate its concentration fluctuations.

For each promoter and each condition, we compared the observed distribution of log GFP concentrations with the distribution that would result from only growth rate fluctuations, i.e., by setting the volumic protein production rate to a constant 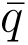 and removing sampling noise at cell division. As shown in Fig. 3, the growth rate fluctuations always contribute a substantial fraction of the total observed expression noise (blue distributions). In addition, as the average growth rate increases with nutrient quality, the concentration distributions resulting from growth rate fluctuations become systematically narrower and account for less of the observed concentration fluctuations of each promoter.

We quantified this behavior by calculating, for each promoter and each condition, the inter-quartile range *η_C_* of log GFP concentration resulting from only growth rate fluctuations (Fig. 4a). First, we find that the protein concentration noise resulting from growth rate fluctuations is the same for all promoters in a given condition. This reflects that the growth rate fluctuations have the same statistics in each experiment in the same growth condition (see also Fig. S2a) and that the variance in log GFP concentration induced by the growth rate fluctuations is independent of the absolute protein production rate 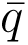 of the promoter. Second, we find that the concentration noise induced by growth rate fluctuations systematically decreases with the mean growth rate.

**Figure 4:**
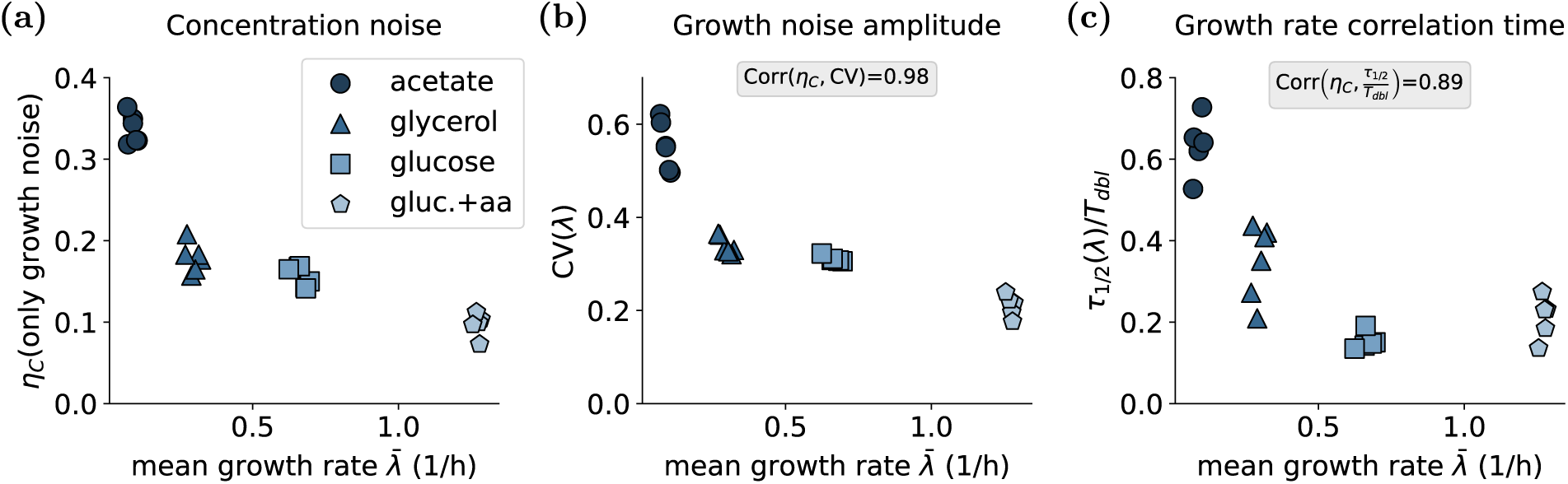
As nutrient quality is varied, growth noise systematically decreases with the average growth rate. (a) Gene expression noise levels *η_C_* resulting from growth rate fluctuations (i.e., inter-quartile ranges of the blue log GFP concentration distributions of Fig. 3) as a function of mean growth rate for each promoter in each condition (different symbols). (b) The growth rate noise amplitude is measured as the coefficient of variation *CV* (*λ*), namely the standard deviation divided by the mean growth rate. (c) The normalized time scale of growth rate fluctuations is measured as the time lag *τ*_1_*_/_*_2_ after which the growth rate auto-correlation drops to 1*/*2 normalized by the mean doubling time *T_dbl_*. Both *CV* (*λ*) and *τ*_1*/*2_*/T_dbl_* decrease with the mean growth rate and correlate strongly with the concentration noise level in (a), as shown in the boxes at the top of (b) and (c).

To understand this systematic behavior, we examined two key properties of growth rate fluctuations in each condition: their amplitudes and the time scale over which fluctuations in growth rate decay. Strikingly, we find that both the amplitude of the growth rate fluctuations and their time scale decrease with growth rate. That is, both the coefficient of variation (CV) of growth rate (Fig. 4b) and the correlation time (Fig. 4c) systematically decrease with average growth rate. Moreover, both the noise amplitude and time scale of the fluctuations highly correlate with the concentration noise (Fig. 4).

The full auto-correlation functions of the growth rate are shown in Fig. S3, again exhibiting slower decay in slow growth conditions even relative to the typical doubling time. Thus, growth rate fluctuations are not only larger but also decay more slowly in slow growth conditions, both driving larger concentration fluctuations than in fast growth conditions. In SI Sec. B, we illustrate the effect of growth rate fluctuations on concentration noise by varying the growth noise amplitude and timescale independently in a mathematical model. Varying either property affects the concentration noise: At high growth noise levels, GFP is diluted at strongly different rates at different time points, and long-lasting fluctuations accumulate the effect of deviations from the mean growth rate over time.

### 2.4 Complex cross-correlation patterns of growth rate and protein production vary across conditions and promoters

Above we saw that there is little correlation between the volumic production rate and growth rate for the synthetic promoter *hi1* when growing on glucose (Fig. 1e). If the volumic protein production and growth rates were fluctuating completely independently, then production and growth rate fluctuations would contribute independently to the concentration noise. However, several observations indicate that this cannot be the case. For example, both growth rate fluctuations (Fig. 4) and the protein production fluctuations (Fig. S2e) decrease with mean growth rate, but the concentration noise as a function of growth rate varies across promoters and often varies non-monotonically as a function of mean growth rate (Fig. S4).

Moreover, while removing the production fluctuations systematically lowers noise levels in almost all conditions, at the slowest growth in acetate removal of protein production noise slightly *increases* the concentration noise. This behavior can be explained by assuming that growth and production fluctuations are positively correlated in this slow growth condition. Indeed, if fluctuations in protein production and growth rate were perfectly correlated, the concentration noise would be zero. We thus decided to systematically quantify the correlations between production and growth rate fluctuations across all promoters and environments.

In particular, for each promoter and condition, we calculated the cross-correlation between the volumic protein production *q*(*t*^′^) and the growth rate *λ*(*t*) as a function of the time difference *t*^′^ − *t* (Fig. 5). We see that, in the slowest growth in acetate, there is a moderate but consistently positive correlation between protein production and growth rate for all promoters, with a maximum at zero time lag (Fig. 5a). This positive correlation explains why the total protein concentration noise in acetate is less than the sum of the contributions of production and growth rate fluctuations, i.e., the production and growth rate fluctuations dampen each other to some extent. However, it should be noted that, because the correlation of protein production and growth rate is only moderate, the amount of dampening in protein concentration fluctuations is a complex and even non-monotonic function of the amplitude of the protein production fluctuations (Supplement Sec. C).

**Figure 5:**
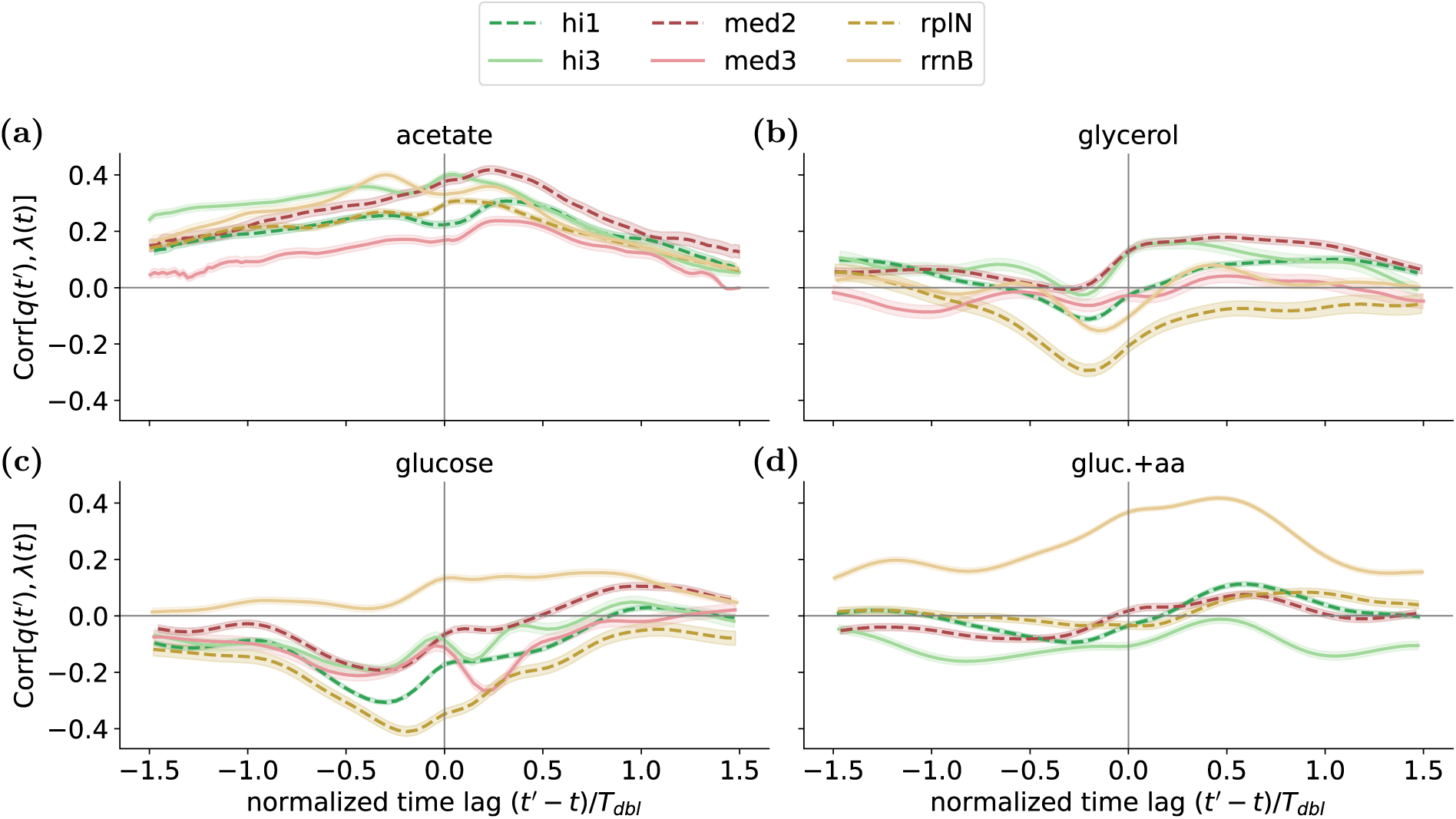
Cross-correlation between growth and volumic protein production varies across promoters and growth conditions. Each panel shows the cross-correlation Corr[*q*(*t*^′^)*, λ*(*t*)] of the production at time *t*^′^ and growth rate at time *t* as a function of the time difference *t*^′^ *t* for each growth condition (panels) and each of the six promoters (see legend). The lines and transparent ribbons show the means of the posterior one standard deviation error. Times on the horizontal axes are normalized by the mean doubling time *T_dbl_* in each condition. Note that, in acetate (a), production and growth are modestly but systematically positively correlated. In the faster growth conditions (c,d), the crosscorrelation functions are more variable across promoters and can also be negative. Note also that the absolute level of the correlations is always modest.

More strikingly, the systematic positive correlation between protein production and growth rate fluctuations that is observed at slow growth in acetate is not observed in any of the other growth conditions, and many promoters exhibit even modest negative crosscorrelations (Fig. 5b-d). This suggests that there must be different mechanisms underlying the protein production fluctuations and that a mechanism that correlates with growth rate dominates protein production fluctuations in acetate, but not in the other growth conditions.

Surprisingly, different promoters exhibit highly distinct cross-correlation patterns of growth rate and production fluctuations. Even synthetic constitutive promoters that express at similar overall levels (similarly colored lines in Fig. 5) show clearly distinct cross-correlation patterns. Moreover, the cross-correlation patterns are quite complex, exhibiting local minima and maxima at different time lags for different promoters. We were quite surprised by the complex cross-correlation patterns with multiple maxima and minima at different time lags that vary across conditions and promoters, and it is tempting to speculate that these might reflect technical artifacts. However, for two promoters (the synthetic promoter (*hi1*) and a ribosomal RNA promoter (*rrnB*) we have data from independent experiments performed at different times using different microfluidic devices, and we find that the cross-correlation curves are remarkably reproducible across these biological replicates, reproducing almost all of the subtle maxima and minima (Fig. S5). This strongly suggests that these patterns reflect real biological correlations in production and growth rate.

Apart from the distinct cross-correlation patterns, different promoters also exhibit clearly different amplitudes of protein production fluctuations (Fig. S2b) that typically decrease with mean growth rate (Fig. S2e). Even the auto-correlation functions of protein production differ not only from condition to condition, but across promoters as well (Fig. S6), with many promoters showing a bi-phasic pattern of an initial fast drop in correlation, followed by a slower decay.

These observations not only suggest that protein production fluctuations result from multiple sources that have different intensities in different conditions, but also that these sources affect different promoters to different extents, even different constitutive promoters.

### 2.5 Binomial sampling at division matters only at low expression

Since, to a first approximation, cytoplasmic proteins are randomly diffusing around the cell, the probability for a given protein to end up in one or the other daughter cell is proportional to the relative sizes of the daughter cells^35–37^. Consequently, the number of proteins ending up in one daughter is approximately a binomial sample of the total protein number and this division sampling noise is often assumed to make a substantial contribution to gene expression noise.

To assess the role of sampling at division for the noise in protein concentrations, we first compared the observed concentration noise with what would be observed if there was no sampling noise at division. That is, we integrate protein concentrations by using the exact time-dependent protein production rates and growth rates, but divide GFP molecules at division in perfect proportion to the sizes of the daughter cells. We find that, for the GFP copy numbers of the transcriptional reporters measured in our experiments, the division noise is negligible (Fig. S7). However, for many genes in *E. coli*, the expression levels are much lower than for the reporters we analyzed here, raising the question of whether division noise becomes a relevant component to the overall concentration fluctuations for lower-expressed genes. To investigate this, we mimic lower-expressed promoters by scaling the observed production rate by a factor *µ_q_*, i.e. we replace the observed volumic production *q*(*t*) with *µ_q_q*(*t*). As a result, the absolute expression levels are rescaled, while growth rate and relative production fluctuations remain unaltered. Moreover, in the absence of cell division noise, the concentration noise level would remain constant, irrespective of the rescaling of the absolute expression level.

We simulate binomial sampling of GFP molecules at cell division and illustrate the procedure in Fig. S8, with the resulting distributions shown in Fig. S9. Figure 6a summarizes our findings on the role of division sampling noise as a function of absolute expression, with the colored symbols showing the measured protein concentration noise for our six promoters, grey symbols showing the concentration noise as the average production is scaled down, and empty symbols showing the concentration noise without sampling noise at division. The results show that sampling noise at division only starts affecting concentration noise when the absolute expression level becomes low.

**Figure 6:**
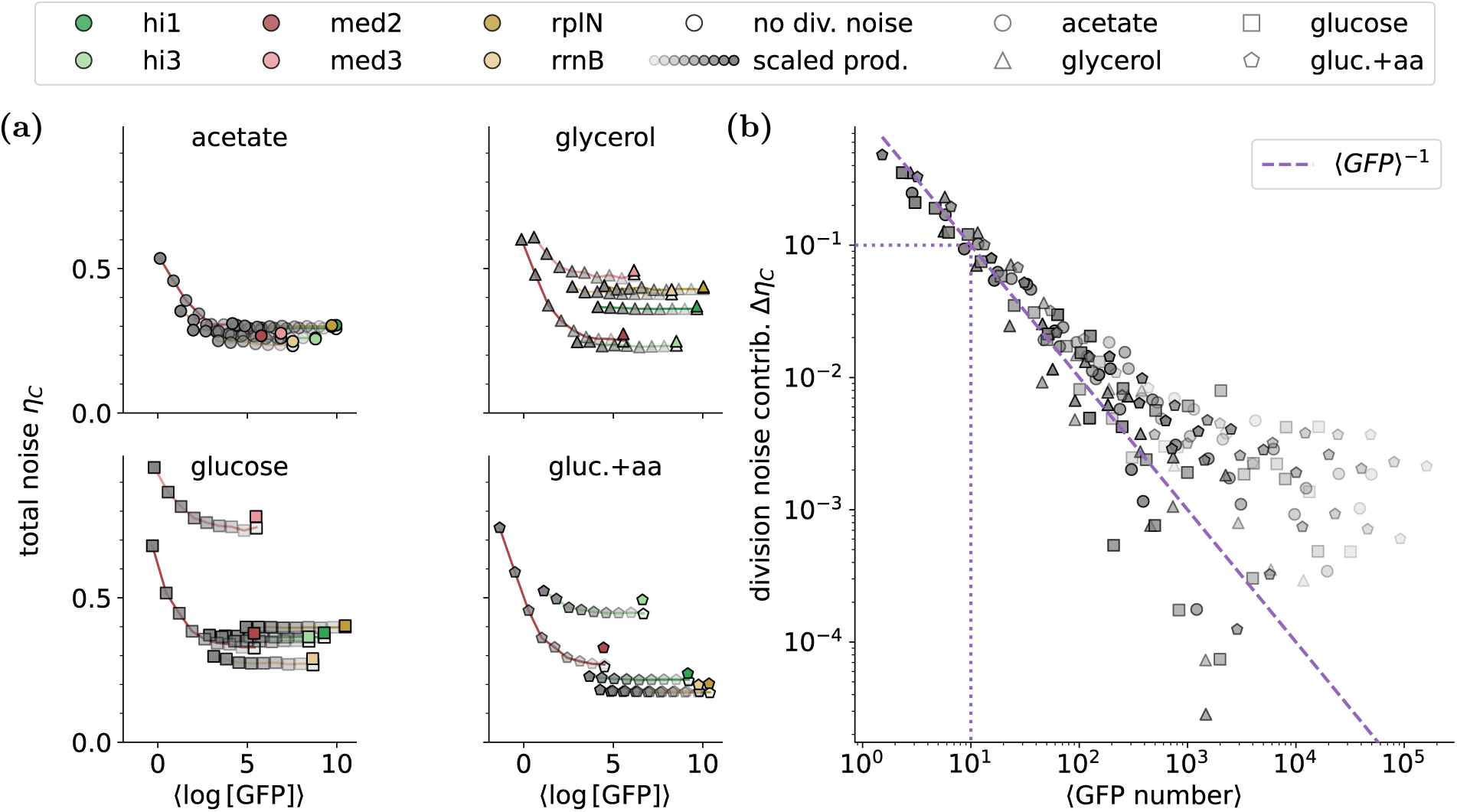
Sampling noise at cell division only becomes important at low expression. (a) Each panel shows the expression noise (inter-quartile range of log protein concentration) for each of our promoters (colored lines) in a given growth condition (indicated at the top) as a function of the mean log-expression level, which is modulated *in silico* by scaling down the observed production rates by an overall factor. The colored symbols correspond to the original data and the grey symbols show the noise level when the production rate is scaled down. As the mean expression level decreases, the relative contribution of division sampling noise increases. Finally, the empty symbols show the case where GFP molecules are partitioned precisely according to the cell sizes of the daughters (no division noise). (b) The noise contribution of the division noise is calculated by comparing the observed noise levels with and without sampling noise at division, Δ*η_C_*= *η_C_*(*µ_q_*) − *η_C_*(no div. noise) as a function of the mean number of GFP molecules per cell, with each dot corresponding to results obtained with rescaled production rates from experiments of a given promoter in a given condition. Note that Δ*η_C_*from different experiments in different conditions all collapse on a common curve that depends only on the mean number of GFP molecules per cell. The dashed line shows the line 1*/x*, which illustrates that the division noise contribution is approximately scaling with the inverse of the mean GFP number. The dotted lines indicate that for 10 GFP molecules per cell, the concentration noise level increases by Δ*η_C_* = 0.1 due to the division noise.

To further quantify how the sampling at division affects concentration noise, we calculated the difference in concentration noise with and without sampling noise at division as a function of the average GFP number per cell for each promoter in each condition (Fig. 6b). The results show that, to a good approximation, the data from all promoters and all conditions collapse onto a common curve. That is, the contribution of sampling at division to protein concentration noise depends *only* on the mean number of GFP molecules per cell and decreases approximately as 1*/*⟨GFP number⟩. Moreover, the concentration noise is only substantially affected at low expression levels, i.e., for a mean expression of 10 proteins per cell, the contribution of the division noise is Δ*η_C_* = 0.1 and, thus, in the absence of other noise sources, the GFP concentration fluctuations are around 10 %.

As discussed in the introduction, at expression levels of 10 or less proteins per cell, the protein production noise is also dominated by the intrinsic noise due to thermal fluctuations. Thus, intrinsic noise and division noise are the dominant noise factors for genes with, on average, less than 10 proteins per cell. But, vice versa, for all genes with substantially higher average expression, neither the intrinsic noise nor the division sampling noise plays a major role in the overall protein concentration noise.

## 3 Discussion

While the functional importance of noise for single cells has been appreciated (see e.g.^9^), the sources and mechanisms responsible for gene expression noise are only partially understood. It is well-appreciated that, due to inherent thermal noise, there are intrinsic fluctuations in the production of both mRNAs and proteins of each gene. However, because the expression variance driven by this intrinsic noise is proportional to the mean, its relative size decreases with expression level and it has been observed^4^ that intrinsic noise is only a major contribution to gene expression noise for genes with 10 or less protein copies per cell on average.

Although, the random partitioning of proteins during cell division is often regarded a potentially substantial source of expression noise, we here found that sampling noise at division is also only a major contributor of gene expression noise for genes with less than 10 proteins per cell on average (Fig. 6). Thus, while intrinsic sampling noise and division noise can be dominant sources of noise for very low expressed genes, these mechanisms do not contribute substantially to expression noise of genes with more than 10 proteins per cell. Gene regulation itself has been shown to be a major source of gene expression noise, with fluctuations in the binding and activities of regulators propagating to their target genes^18,22^. Consequently, gene expression noise is highly condition-dependent, more regulated genes tend to be more noisy, and constitutive (i.e. unregulated) genes are the least noisy in each condition.

Here we focused on understanding how fluctuations in protein production and growth determine protein concentration fluctuations of constitutive and ribosomal genes across growth conditions. Although growth rate fluctuations are typically overlooked as a potential source of gene expression noise, we find that, by causing fluctuations in dilution rate, growth rate fluctuations are a major source of protein concentration fluctuations, on par with protein production fluctuations. Moreover, when comparing growth rate fluctuations across growth conditions with different nutrients, we find that growth rate fluctuations systematically increase as average growth rate decreases, both in their amplitude and in their duration.

Why do growth rate fluctuations increase in slower growth? Very roughly speaking, cells grow slower in poorer nutrient conditions because they need to invest gene expression in a wider array of enzymes to run a larger number of reactions in order to produce the basic building blocks, e.g. amino acids, nucleic acids, and lipids, from the nutrients available in the environment. It is tempting to speculate that growth rate fluctuations are larger and longer lasting at slow growth because there is simply a larger number of steps and components in the metabolic network that may cause bottle necks when expression levels are unbalanced. But independent of this hypothesis, since the growth rate is a compound phenotypic trait, the larger growth rate fluctuations support that phenotypic noise is higher in slow-growth conditions. This is in line with previous studies that found gene expression noise generally decreases with growth rate^22^.

Since we found that growth rate fluctuations are a major driver of gene expression noise, it is also tempting to speculate that the decrease of gene expression noise with average growth rate is caused by the decrease of growth rate fluctuations. If this interpretation is correct, it would constitute an interesting self-reinforcing mechanism driving phenotypic heterogeneity: At slow growth there are more components whose expression fluctuations can cause bottlenecks, which leads to larger and longer lasting growth rate fluctuations, which in turn drive larger gene expression fluctuations.

In contrast to growth rate fluctuations, production fluctuations do not show any consistent changes with the average growth rate in different conditions. In most growth conditions, there are only weak cross-correlations between growth and protein production fluctuations, and the sign and form of the cross-correlation varies across promoters. Only in the slowest growth in acetate do we find that there is modest, but systematically positive cross-correlation between growth and production. We speculate that in this slow growth condition, growth rate fluctuations may be driven to a substantial extent by variation in overall translation activity, so that when growth rates are higher, the protein production of all genes is higher, explaining the systematic positive correlation. Vice versa, the lack of systematic correlations between growth and production in the other conditions suggest that growth rate fluctuations are driven by something other than variation in translation activity. It is conceivable that growth rate fluctuations are driven by fluctuations in the total mRNA concentration in these conditions.

Surprisingly, despite lacking known regulatory input and identical positions on the chromosome, the correlation functions differ between the four synthetic constitutive promoters we measured. Moreover, the production strongly varies between promoters with distinct behaviors between promoters across growth conditions. As a result of the non-trivial correlation structure and production noise behavior, the overall concentration noise of individual promoters changes non-monotonically with growth rate and is highly promoter-dependent.

These observations raise the question of what mechanisms might be involved in the complex fluctuations in protein production rates. First, in a recent analysis of analogous single-cell data, we observed that there are systematic changes in growth rate and production across the cell cycle, with different promoters exhibiting different cell cycle dependence in production^25^. Thus, some of the observed production fluctuations are associated with fluctuations along the cell cycle, although these cell cycle dependencies are generally small, and the mechanistic origins of these cell cycle dependencies are also not understood.

It is challenging to hypothesize what mechanisms of production rate fluctuations could affect different promoters so differently. Notably, since the only difference between our reporter constructs is in the promoter, i.e., the GFP mRNAs produced are essentially the same, it is unlikely that these differences result from differences in either translation rates or mRNA stability. That is, the promoter-dependent patterns of production fluctuations and their cross-correlation with growth rate fluctuations must derive predominantly from differences in transcription.

It is noteworthy that the promoter with by far the largest amplitude in production fluctuations is *med3*, and in recent work we have established that this promoter is predominantly transcribed by the rpoS sigma factor^38^. If each promoter is transcribed to different extents by different sigma factors, and these sigma factors exhibit different concentration fluctuations in different environments, then this might explain some of the promoter-dependent fluctuations in protein production.

A final noteworthy observation is that the only promoter that exhibits positive crosscorrelation with growth rate at fast growth is the ribosomal RNA promoter *rrnB*. If growth rate fluctuations at fast growth are driven by fluctuations in total mRNA concentration, then it would be conceivable that the ribosomal RNA promoter (*rrnB*) would exhibit the highest correlation in its expression with the total mRNA level.

In conclusion, our work highlights the importance of growth rate fluctuations in driving phenotypic heterogeneity of single cells and finds that growth rate fluctuations decrease as the average growth rate increases, potentially explaining why phenotypic heterogeneity is generally lower at fast growth. In contrast, protein production fluctuations are surprisingly variable across promoters, differing even between synthetic constitutive promoters. This suggests that, in contrast to growth rate fluctuations, protein production fluctuations are driven by multiple mechanisms that have different intensity in different growth conditions and affect different promoters to different extents. Uncovering what these mechanisms are, and why they affect even different constitutive promoters to different extents, is an important topic for future studies.

## 4 Methods

### Mother Machine experiments

We used 6 strains that each have promoter-GFPmut2 cassettes integrated at the HKO22 locus attB of the chromosome of the *E. coli* K-12 MG1655 strain. The synthetic promoter sequences, i.e., *hi1*, *hi3*, *med2*, and *med3* were taken from Ref.^18^. Ribosomal promoters, namely *rplN* and *rrnB* were derived from the Zaslaver library^39^. All promoter sequences are provided in the Supplementary Materials Sec. D. Cells were grown in the dual-input Mother Machine described in Ref.^24^ with M9 minimal media supplemented with different nutrients. This includes 0.05 % acetate, 0.4 % glycerol, 0.2 % glucose, and 0.2 % glucose plus with amino acids. All experiments were conducted at 37 ^◦^C. Images were acquired every 12 min, 6 min, 3 min or 1.5 min for cells grown on acetate, glycerol, glucose, and glucose+aa, respectively.

### Image analysis

We analyzed the microscopy data with a new version of the image analysis software MoMA introduced in Ref.^24^. The new version uses a U-Net to segment cells. The tracking of cells is based on a cost function as described previously, which has been adjusted as necessary.

### Cell dynamics inference

We denoised the Mother Machine data sets using the inference method RealTrace^25^. We obtain posterior distributions for cell log size *x*(*t*), GFP content *g*(*t*), growth rate *λ*(*t*), and volumic production rate *q*(*t*) at each measurement time point. Two-time point posterior distributions were used to calculate auto and cross-correlation functions as described in Ref.^25^. The bleaching rate was estimated from separate experiments with values *β* = 0.048, 0.096, 0.192, 0.384 h^−1^ for acetate, glycerol, glucose, and glucose+aa, respectively.

The traces of the total GFP number *g* and volumic production *q* are corrected for autofluorescence using *g*(*t*) → *g*(*t*) − *c*_autofl._*l*(*t*) and *q*(*t*) → *q*(*t*) − *c*_autofl_.(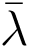+ *β*), respectively.

Here, *c*_autofl._ is the auto-fluorescence per cell length determined from separate experiments (see Tab. 1) and 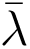 the average growth rate in the respective growth condition.

### Numerical integration

During the cell cycle, the dynamics of the total GFP content *g* reads,

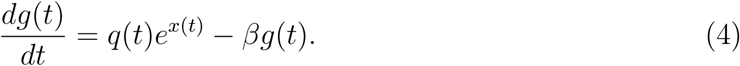

For the numerical forward integration with a fixed volumic production rate *q*(*t*) = ⟨*q*⟩, each step from *t* to *t* + *dt* is calculated according to,

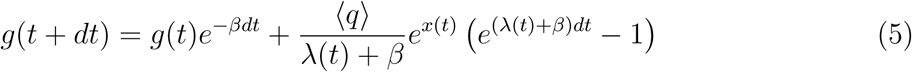

Here, we used the approximation of a constant *λ*(*t*) for the time interval (*t, t* + *dt*).

For simulations that include binomial sampling at cell division, the GFP content of one daughter *g*(*t*+*dt*) cell is drawn from a binomial distribution. Each of the *g*(*t*) GFP molecules of the mother cell has a probability of being found in the daughter cell after the division, which is given by the length of the daughter cell divided by the length of the mother cell,

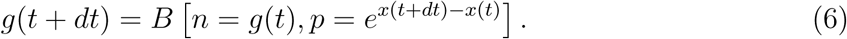

The second daughter cell, if present, obtains the remaining GFP content.

## Data and code availability

All experimental data sets are available at https://doi.org/10.5281/zenodo.18129208. The code for the analysis and visualization is available at https://github.com/bjks/ fluctuations_paper_analysis.

## Acknowledgments

We would like to thank Bor Kavčič and Daan de Groot for helpful discussions and feedback. This research was supported by the Swiss National Science Foundation’s grants numbers 159673, 184937, and Sinergia grant number 189910.

## Supplementary Materials

## A Supplementary Figures

**Figure S1:**
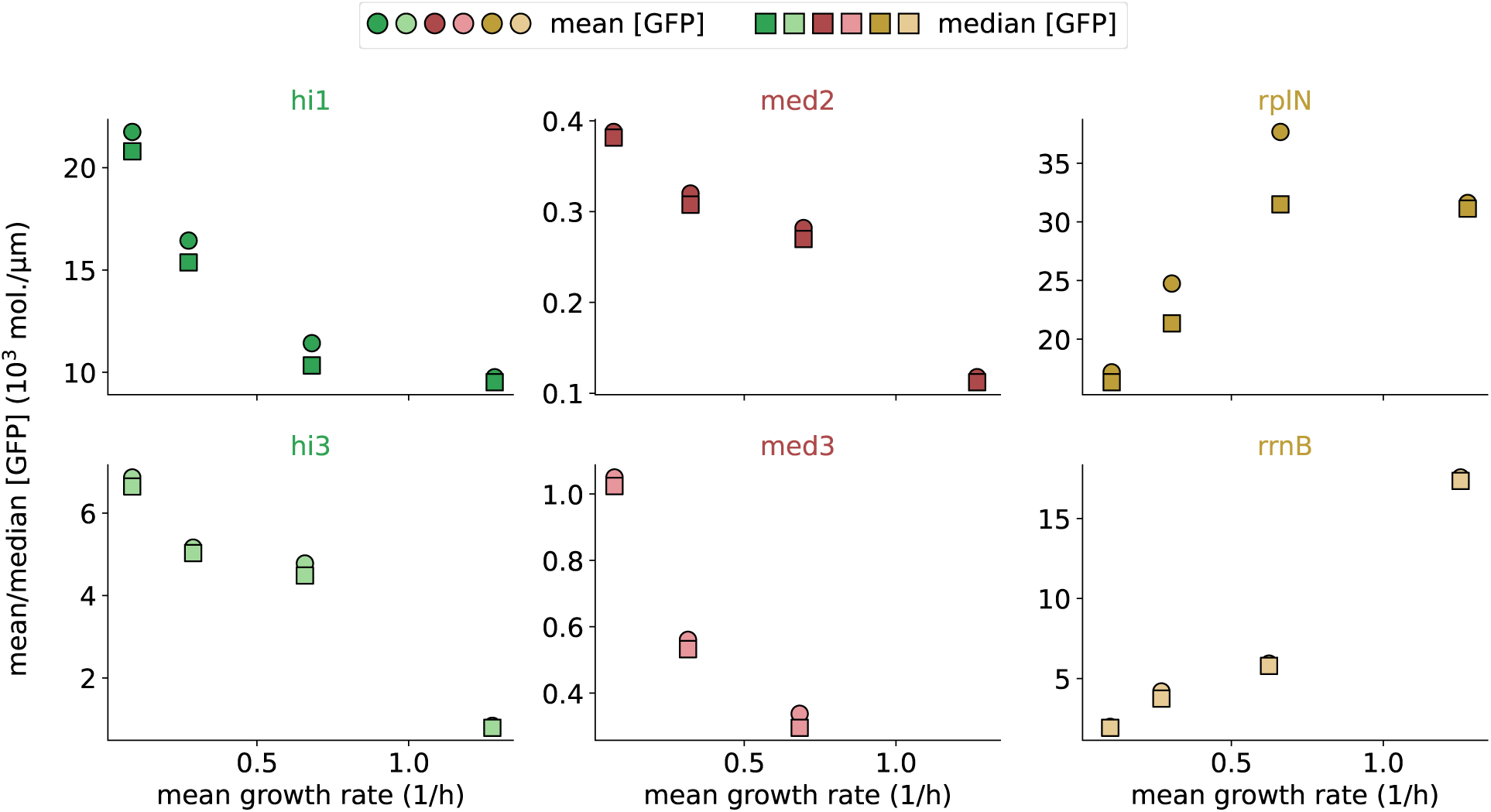
Mean and median concentration across growth conditions. The mean (circles) and median (squares) concentrations as a function of the mean growth rate across growth conditions. The dots from left to right correspond to acetate, glycerol, glucose, and glucose+aa. The concentration of all synthetic, constitutive promoters (*hi1*, *hi3*, *med3*, and *med3*) decreases with the mean growth rate. In contrast, the concentration of ribosomal promoters (*rplN* and *rrnB*) typically increases with the mean growth rate. The concentration distribution of *rplN* in glucose is very long-tailed. As a result, the median concentration in this case is notably lower than the mean. The median concentration of *rplN* does not further increase from glucose to glucose+aa.

**Figure S2:**
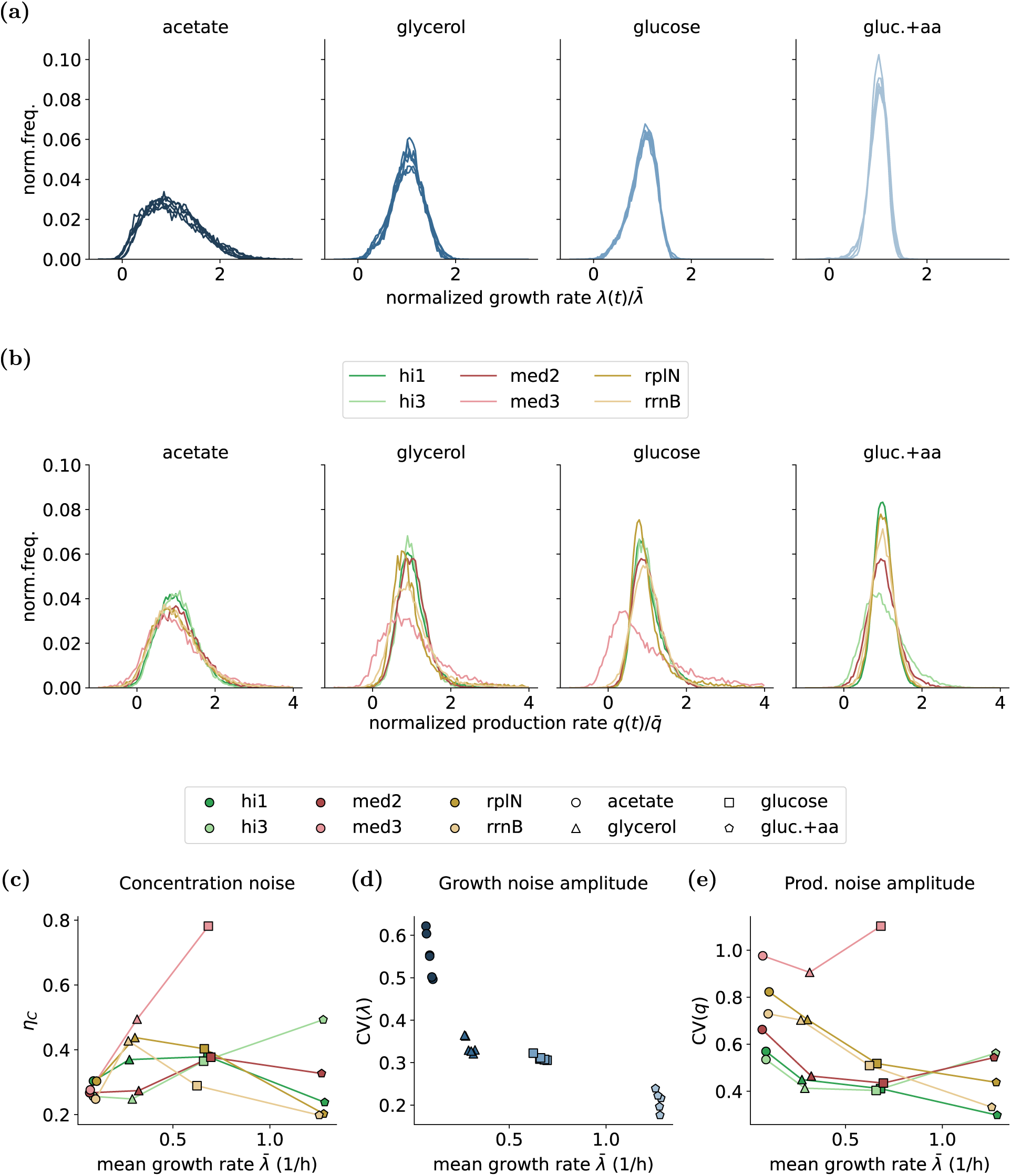
Concentration, growth, and production noise amplitudes exhibit different dependences on the mean growth rate. (a) Distributions of the instantaneous growth rates normalized by the mean growth rate. The distributions are very reproducible across data sets and become narrower in faster growth conditions. (b) Distributions of the instantaneous volumic production rates normalized by the mean rate. The distributions of normalized production rates are promoter dependent. (c) Noise level of the GFP concentration (interquartile range of the log concentration) measured for different promoters and growth conditions. (d) Growth rate noise amplitude measured as the CV (coefficient of variation), replotted from Fig. 4(d). The CV corresponds to the widths of the distributions of normalized growth rates in (a). (e) Volumic production rate noise amplitude tends to decrease with the mean growth rate. However, some promoters, notably *med3* in glucose (light red square), increase production noise at a faster growth rate. Together, this shows that the total concentration noise is not explained by independent contributions of growth and production fluctuations.

**Figure S3:**
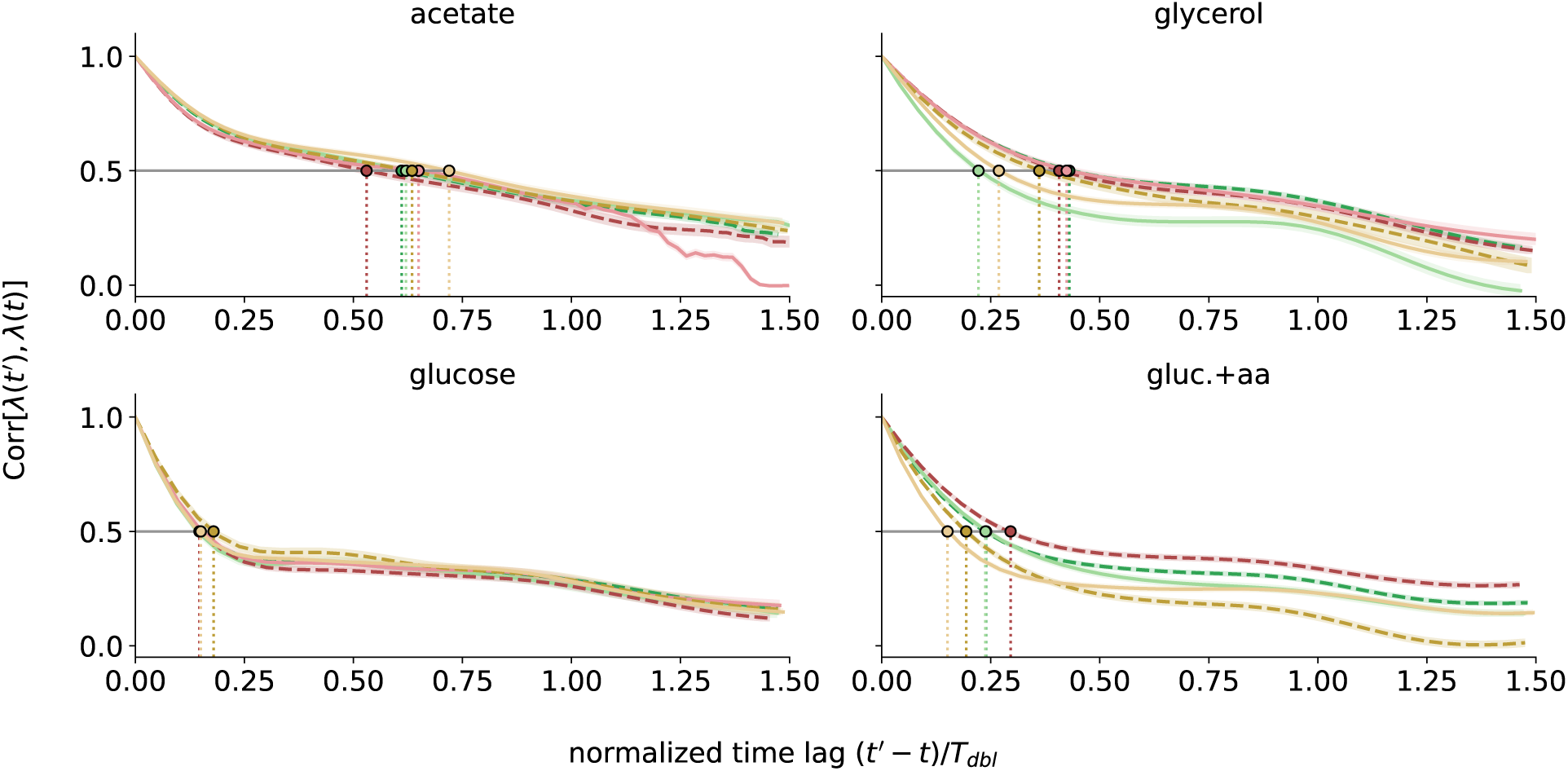
Auto-correlation functions of the growth rate. *λ*(*t*) **across conditions.** Each panel shown the auto-correlation function of growth rate from experiment with different promoters (colors). The transparent ribbon show two standard-deviation error bars. The time lag on the x-axis is normalized with the respective mean doubling time in each condition. Thus, the x-axis can be read as cell cycle fractions. The vertical dotted lines indicate the time lags at which the correlation function drops to 0.5. The normalized time lag that is determined in this way is plotted in Fig. 4(c), showing that fluctuations have longer durations in slow growth conditions, even relative to the respective cell cycle time.

**Figure S4:**
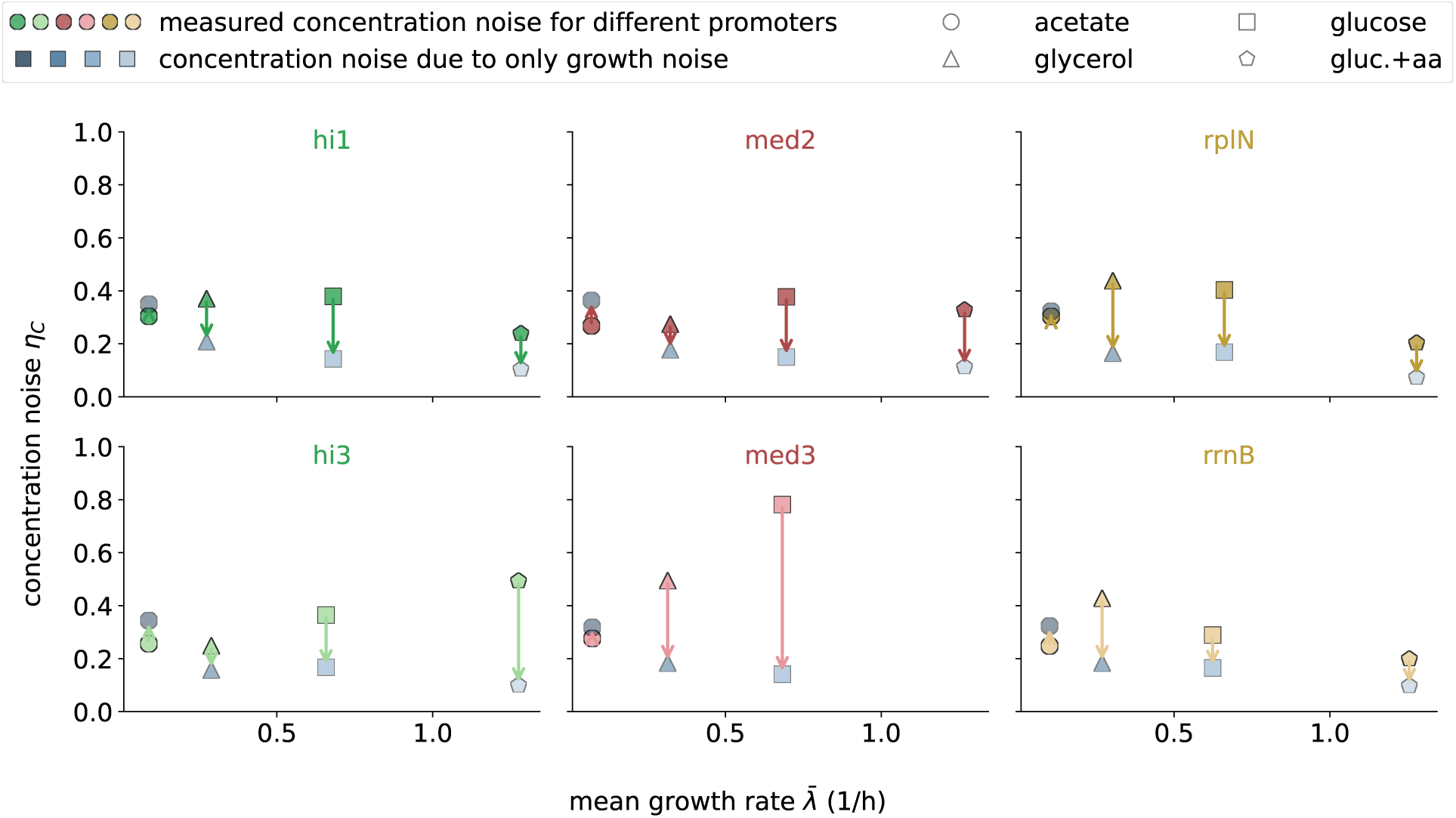
The effect of growth rate fluctuations on concentration noise varies across promoters and growth conditions. Each panel shows noise levels (inter-quartile range of log GFP concentration) as a function of average growth rate) for one promoter (indicated at the top). The concentration noise levels as measured experimentally are shown as colored symbols, and the concentration noise levels predicted when the concentration noise is solely driven by growth rate fluctuations are shown as blue symbols. Note that the latter are always lower than the measured noise levels, except for in acetate. That is, in acetate the growth and production fluctuations dampen each other to some extent, so that when production fluctuations are removed, the noise increases.

**Figure S5:**
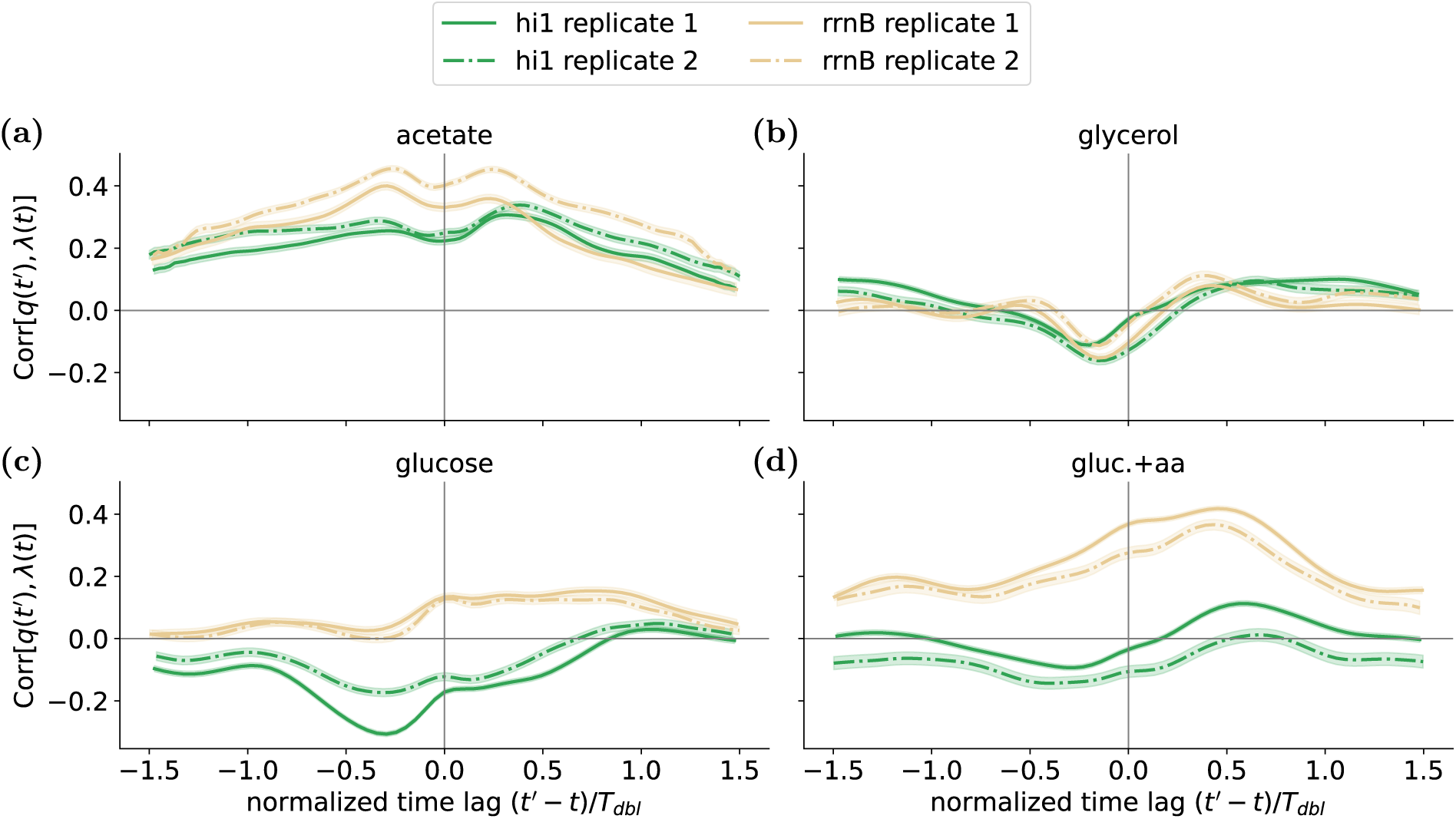
The cross-correlation of growth and production is highly reproducible across replicate experiments. The cross-correlation functions between the volumic production *q* and the growth rate *λ* for the synthetic constitutive promoter *hi1* (green) and the ribosomal RNA promoter *rrnB* (beige) across four growth conditions (panels). The solid and dotted lines correspond to results from two biological replicate experiments using different microfluidic devices on different dates. Note that even subtle local maxima and minima in the cross-correlation functions are highly consistent across the replicates.

**Figure S6:**
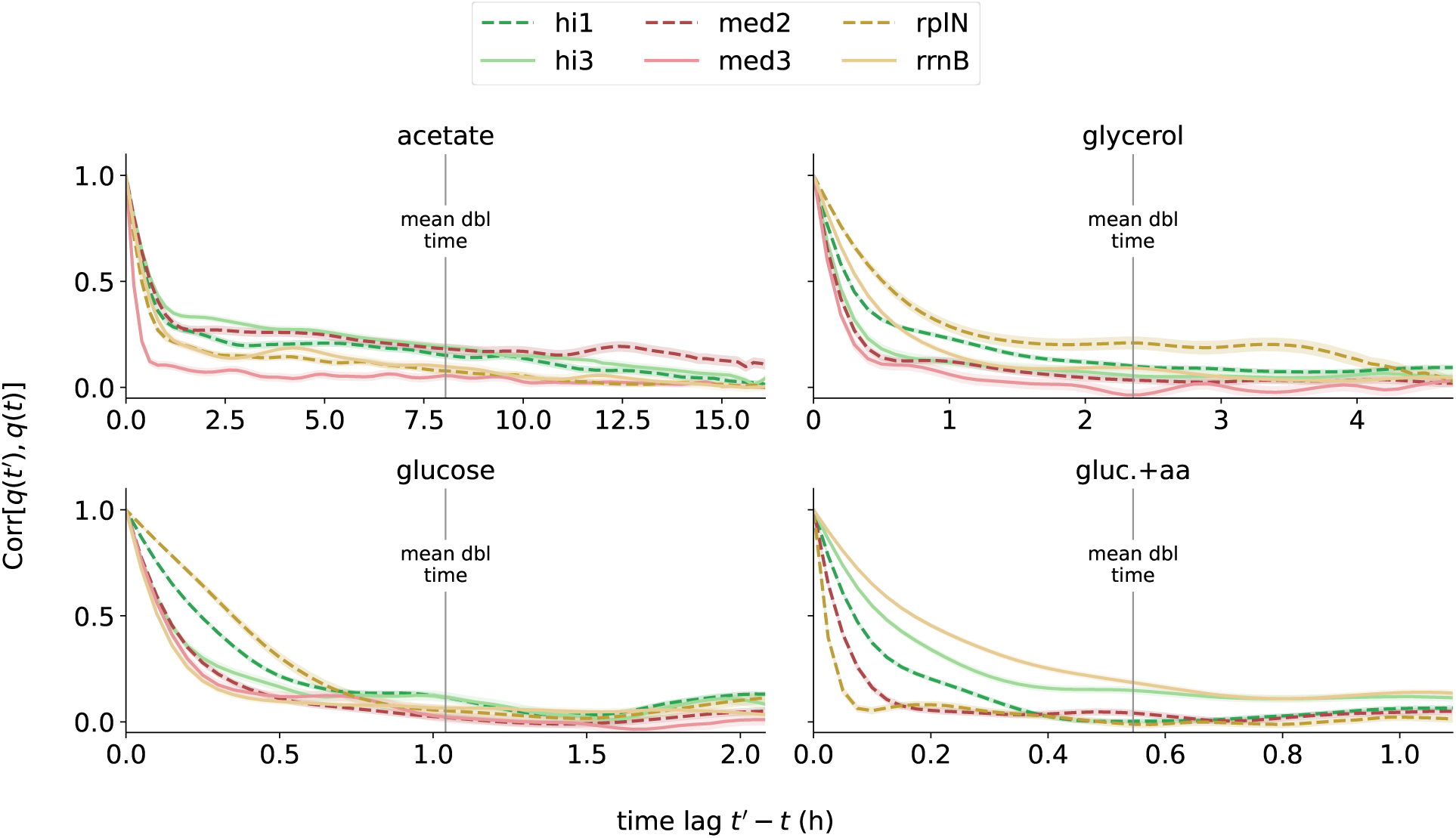
Auto-correlation functions of volumic production rate. *q*(*t*) **across promoters and growth conditions.** Each panel shows the auto-correlation functions of volumic production as a function of time for the 6 different promoters (colored solid and dashed lines, see legend) in one growth condition (indicated at the top). The production auto-correlation functions vary across promoters and growth conditions. The transparent ribbons show two standard-deviation error bars. As a guide for the eye, the mean doubling time (averaged over all data sets in the respective condition) is indicated by a vertical line in each panel.

**Figure S7:**
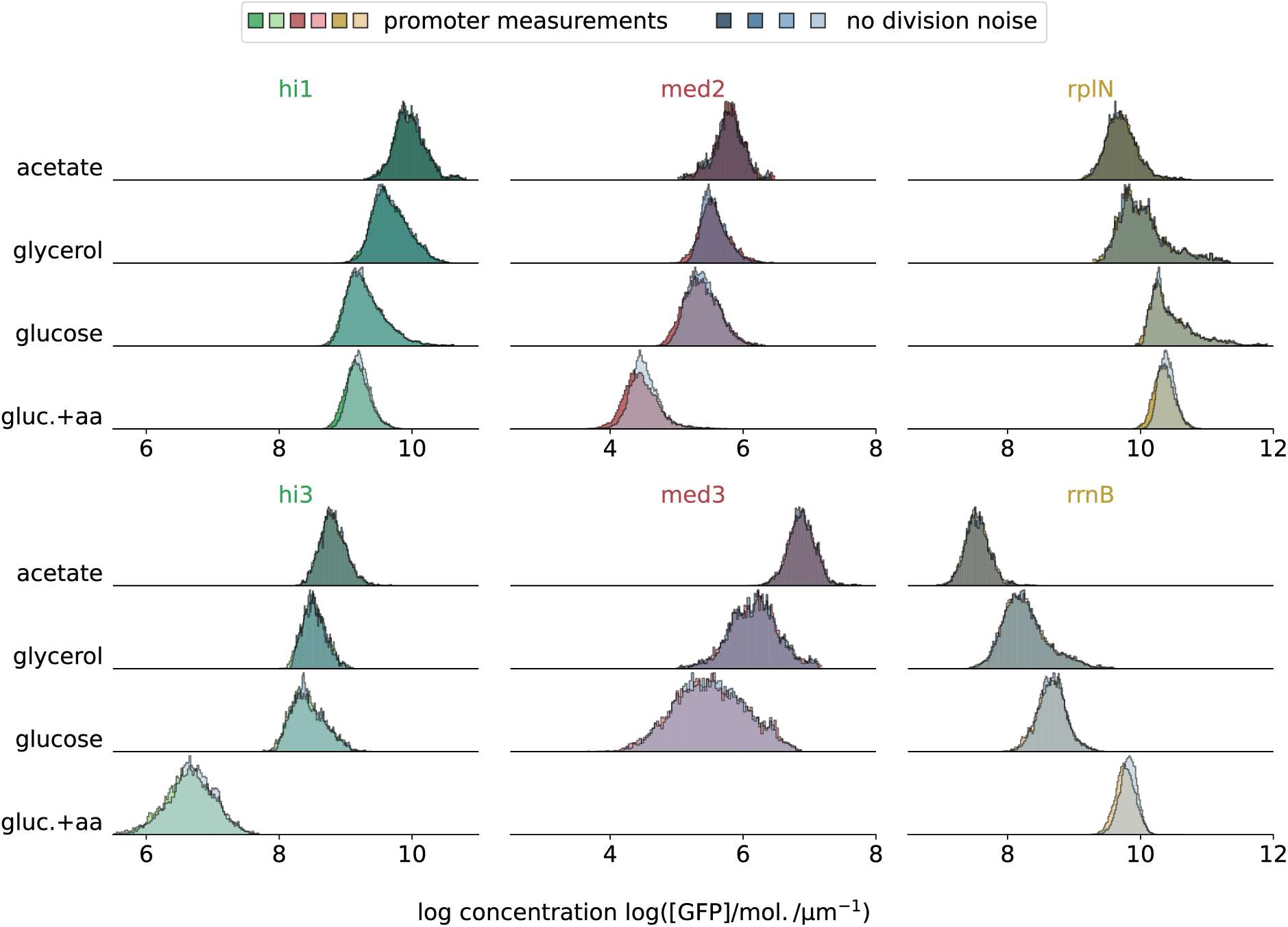
Impact of division noise is negligible for the reporters measured in our experiments. Histograms of log GFP concentration as measured experimentally (colors) and when the division noise is eliminated and GFP molecules are precisely distributed according to the sizes of the daughter cells (grey). Each panel corresponds to a promoter (indicated at the top) with different rows corresponding to different growth conditions (indicated on the left). Note that the grey histograms, which represent the dynamics without division sampling noise, are almost identical to the measured distributions, implying that the division noise is negligible for the GFP concentration noise of these transcriptional reporters.

**Figure S8:**
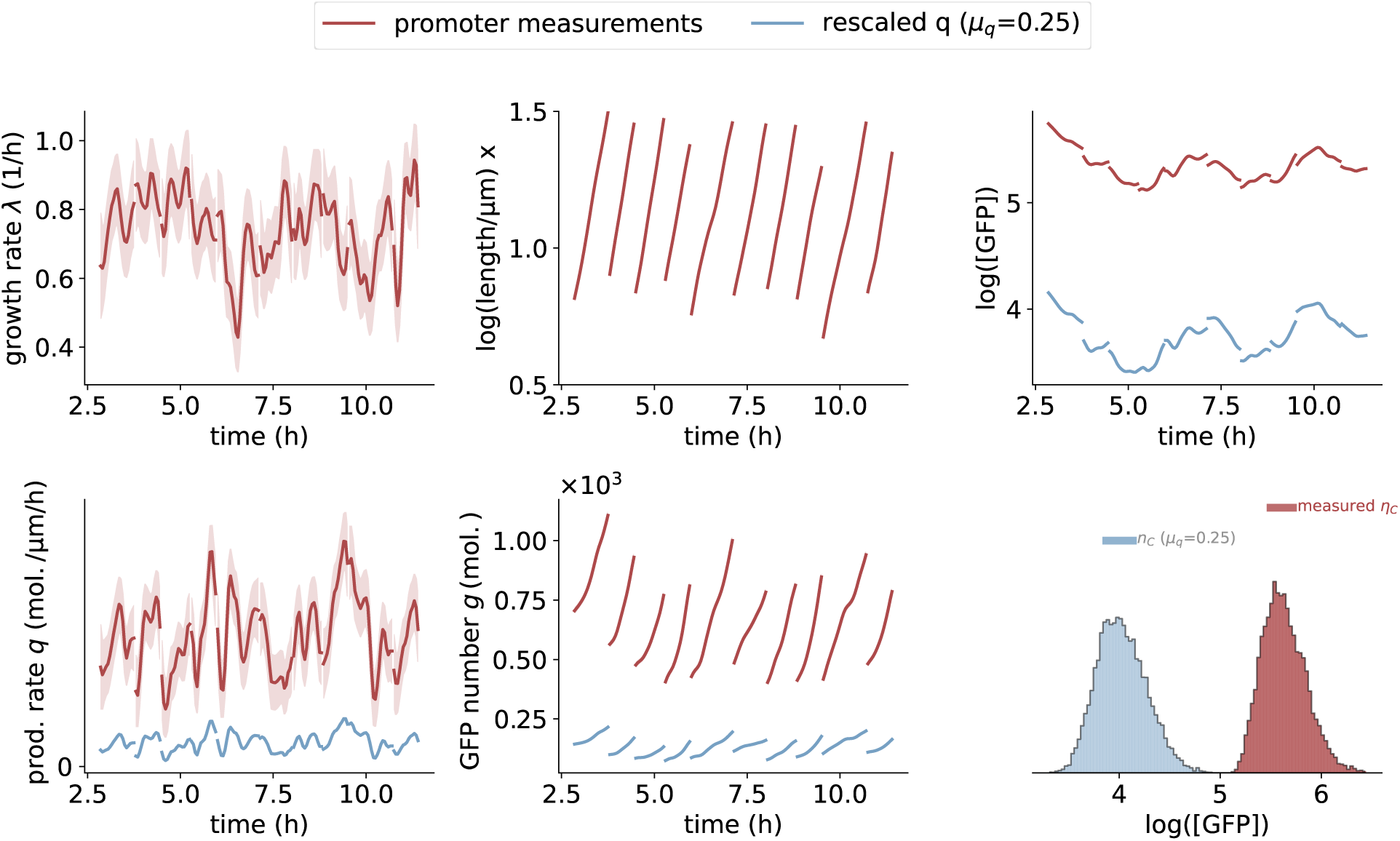
Systematically scaling the volumic production rate to alter the mean expression level. By scaling the production by a factor *µ_q_*, i.e., *q*(*t*) → *µ_q_q*(*t*), we mimic the expression dynamics of a promoter with lower average expression levels, allowing us to estimate the impact of the binomial sampling at division, without altering any of the other features of the expression and growth dynamics. Shown is an example lineage of cells growing on glucose expressing GFP under the control of the *med2* promoter. Panels show, in lexical order, the growth rate, log-cell size, log GFP concentration, volumic production rate, GFP molecules per cell and histograms of log-GFP concentration for the original measurements (red) and dynamics when the production rate is scaled by a factor *µ_q_* = 0.25. Note that the down-scaling of absolute expression increases the width of the concentration histogram, illustrating that sampling at division plays an increased role at lower expression levels.

**Figure S9:**
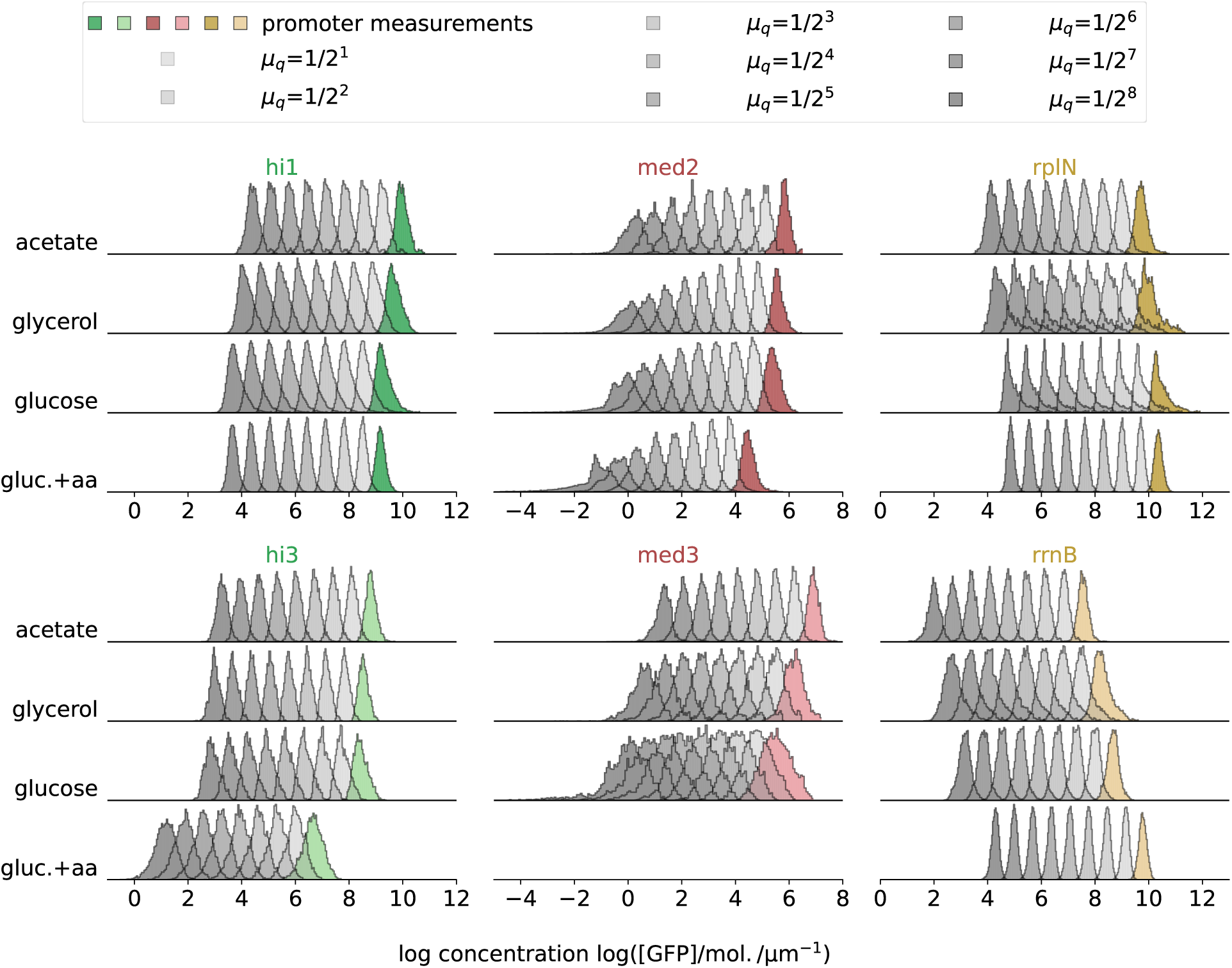
Log GFP concentration distributions at different expression levels. By mimicking expression and growth dynamics at different absolute expression levels, we assess the impact of partitioning noise at cell divisions. The colored distributions correspond to the experimentally measured dynamics. Notably, these distributions are almost entirely driven by growth and production fluctuations, and the division noise is negligible. The grey distributions correspond to results from identical dynamics apart from the fact that the production rates have been scaled down by different amounts (see legend). The noise levels of these distributions increase due to the increased relative contribution of partitioning noise at cell divisions. This is especially noticeable for promoters that are already lowly expressed. The resulting concentration noise levels are shown in Fig. 6.

**Table 1:**
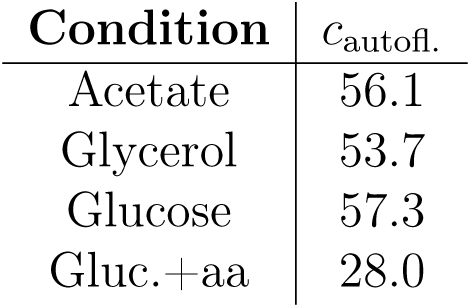
Auto-fluorescence correction factor. *c*_autofl._, the auto-fluorescence per cell length, was determined by measuring the fluorescence of strains without GFP in the Mother machine. *c*_autofl._ is given in units that correspond to numbers of GFP molecules per micron.

| Condition | $c_{\text{autofl.}}$ |
| --- | --- |
| Acetate | 56.1 |
| Glycerol | 53.7 |
| Glucose | 57.3 |
| Gluc.+aa | 28.0 |

## B Impact of growth rate fluctuation magnitude and time scale on concentration noise

We explore the impact of growth rate fluctuations on the concentration noise as the magnitude and the time scale of growth rate fluctuations are varied. For that, we formulate a toy model for stochastic growth that allows us to independently vary both properties.

We use an Ornstein-Uhlenbeck process to model the instantaneous growth rate,

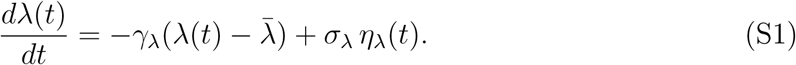

The Ornstein-Uhlenbeck process is characterized by a mean growth rate 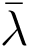, an inverse correlation time scale *γ_λ_*, and a characteristic kick size *σ_λ_*. *η_λ_*(*t*) denotes white noise with mean 0 and standard deviation 1. The mean growth rate is set to the mean growth rate of cells growing on glucose 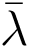= 0.69 1*/*h (glucose *hi1* data set). The parameters *γ_λ_* and *σ_λ_* are set by varying the normalized correlation time *τ*_1*/*2_*/T_bl_* and the noise level *CV* (*λ*), which are related to the parameters via

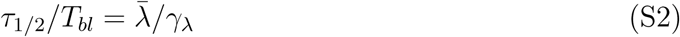

and

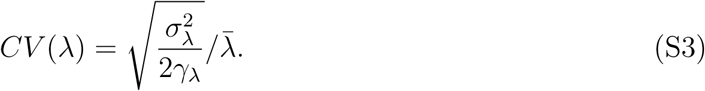

These quantities are varied to cover the experimentally found range (see Fig. 4).

We simulate 50 lineages of 100 cell cycles after discarding 5 cell cycles of each lineage and determine the concentration noise level. Fig. S10 shows the concentration noise as a function of the normalized correlation time *τ*_1*/*2_*/T_bl_* and the noise level *CV* (*λ*). Increasing the normalized correlation time and the noise level both increase the concentration noise.

**Figure S10:**
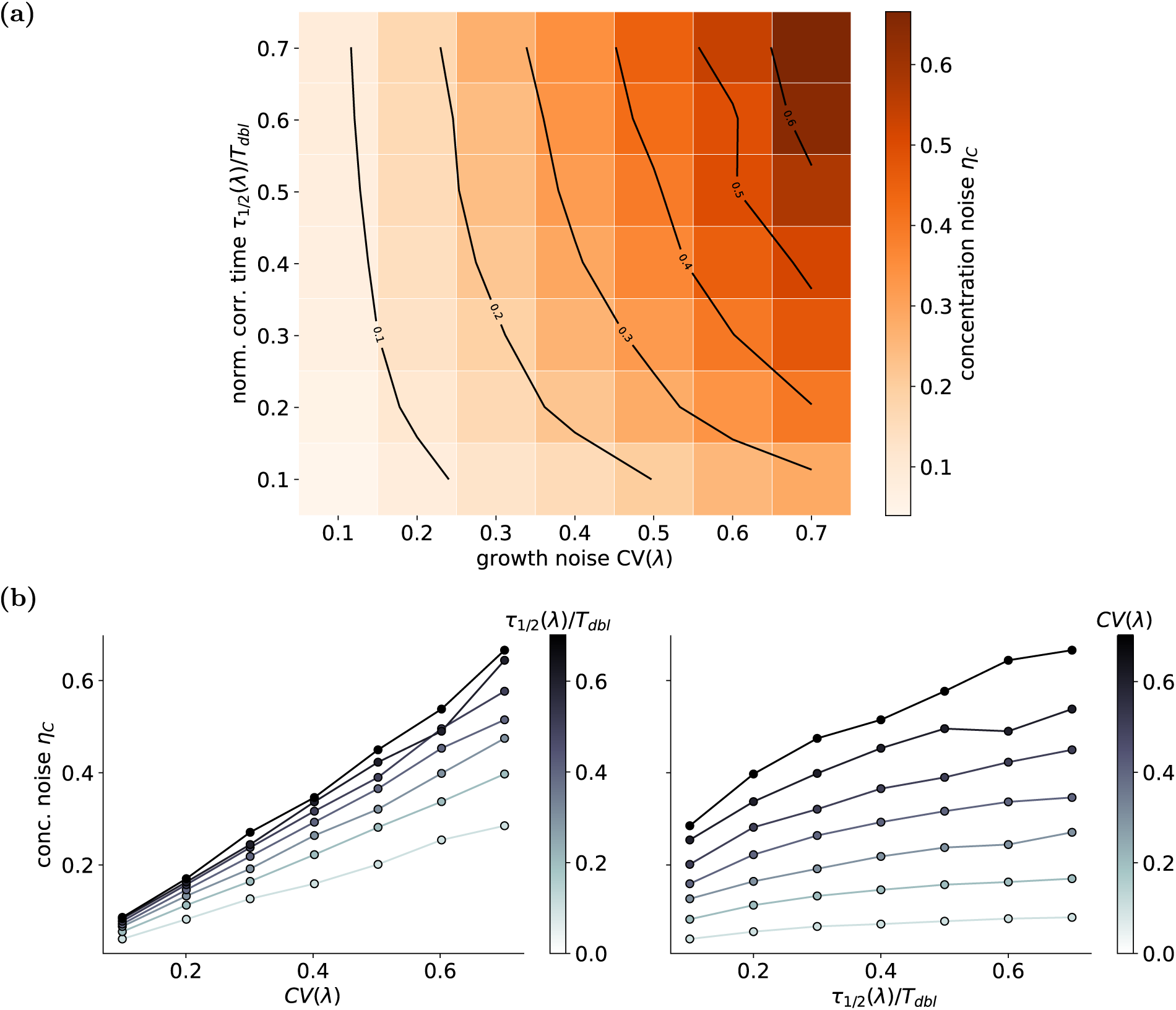
Concentration noise as a function of growth noise. *CV* **and normalized time scale of the growth fluctuations** *τ*_1*/*2_(*λ*)*/T_dbl_*. (a) Concentration noise increases with the *CV* (*λ*) and *τ*_1*/*2_(*λ*)*/T_dbl_*. (b) Replotted concentration noise from (a) as a function of the growth noise level *CV* (*λ*) for different time scales *τ*_1*/*2_(*λ*)*/T_dbl_* and *vice versa*.

## C Impact of growth-production coupling

The positive correlation between the growth rate and production rate of cells growing in acetate results in GFP concentration noise levels that are slightly lower than the noise levels predicted for cells with a constant volumic production rate. Thus, the observed production fluctuations dampen the growth rate fluctuations, which hints at a non-monotonic dependence of the concentration noise level on the noise level of the production in this condition.

We explore this further by altering the noise level of the production *in silico*: By scaling the production by a factor *α_q_*, i.e. *q*(*t*) = ⟨*q*⟩ + *δq*(*t*) → ⟨*q*⟩ + *α_q_δq*(*t*) we can alter its noise level without affecting the cross-correlation function of growth and production. Example distributions for the promoters *med2* and *rrnB* are shown in Fig. S12(a) and (b) and the noise level of the GFP concentrations across promoters is shown in Fig. S12(c). The positive growth-production correlation functions in acetate result in concentration noise levels for low *CV* (*q*) that are slightly below the production noise-free cases (Fig. S12(d)). The measured production noise levels lie in the range of *CV* (*q*) to indeed decrease the concentration noise. This can also be observed for *rrnB* in glucose+aa. However, the measured *CV* (*q*) in this case is outside the range of production noise levels that would lower the IRQs.

**Figure S11:**
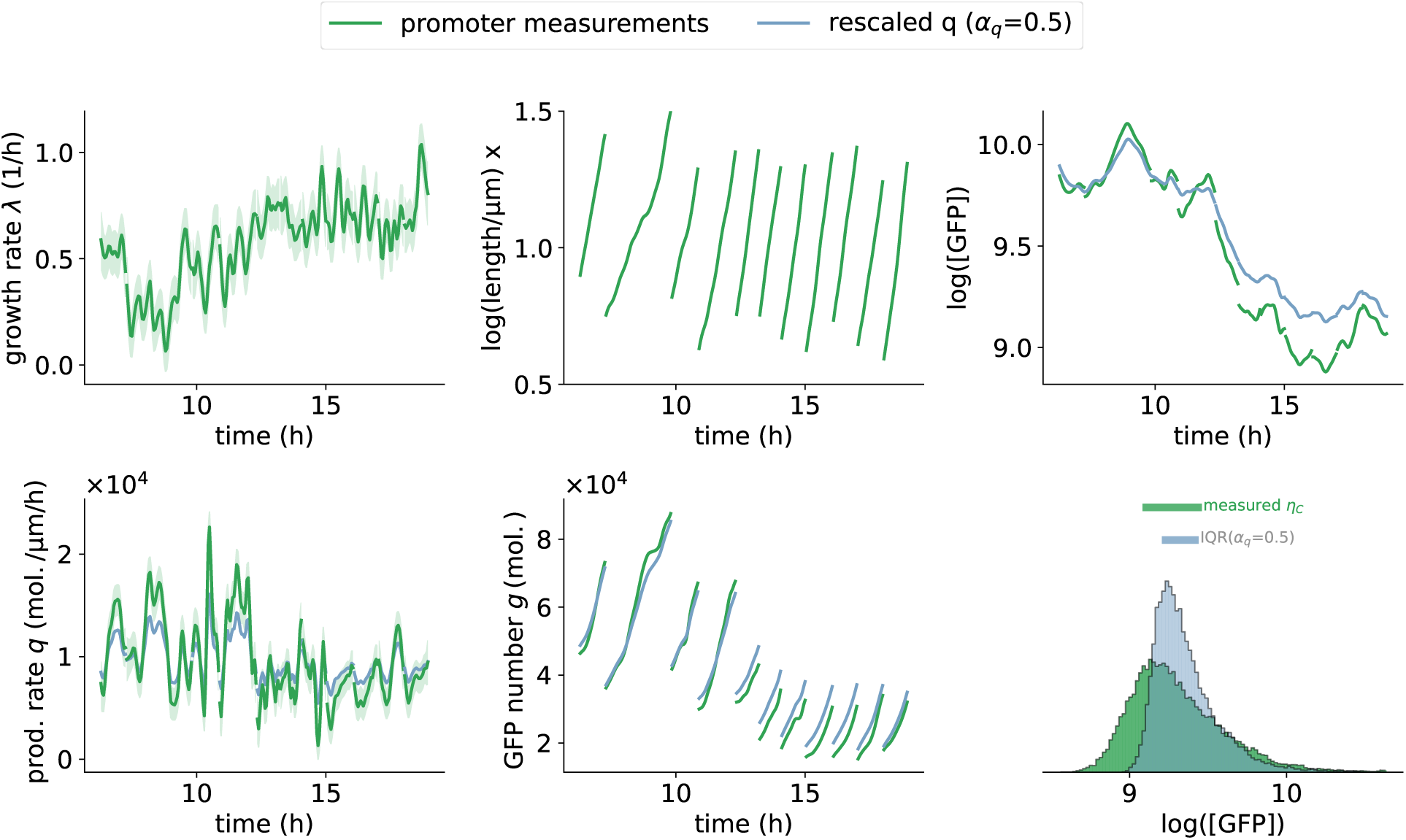
Scaling of the production noise level. *CV* (*q*). Example lineage of cells growing on glucose for the promoter *hi1*. In green is the cell dynamics as experimentally measured, and in blue is the cell dynamics when the production fluctuations are scaled according to *q*(*t*) → ⟨*q*⟩ + *α_q_δq*(*t*).

**Figure S12:**
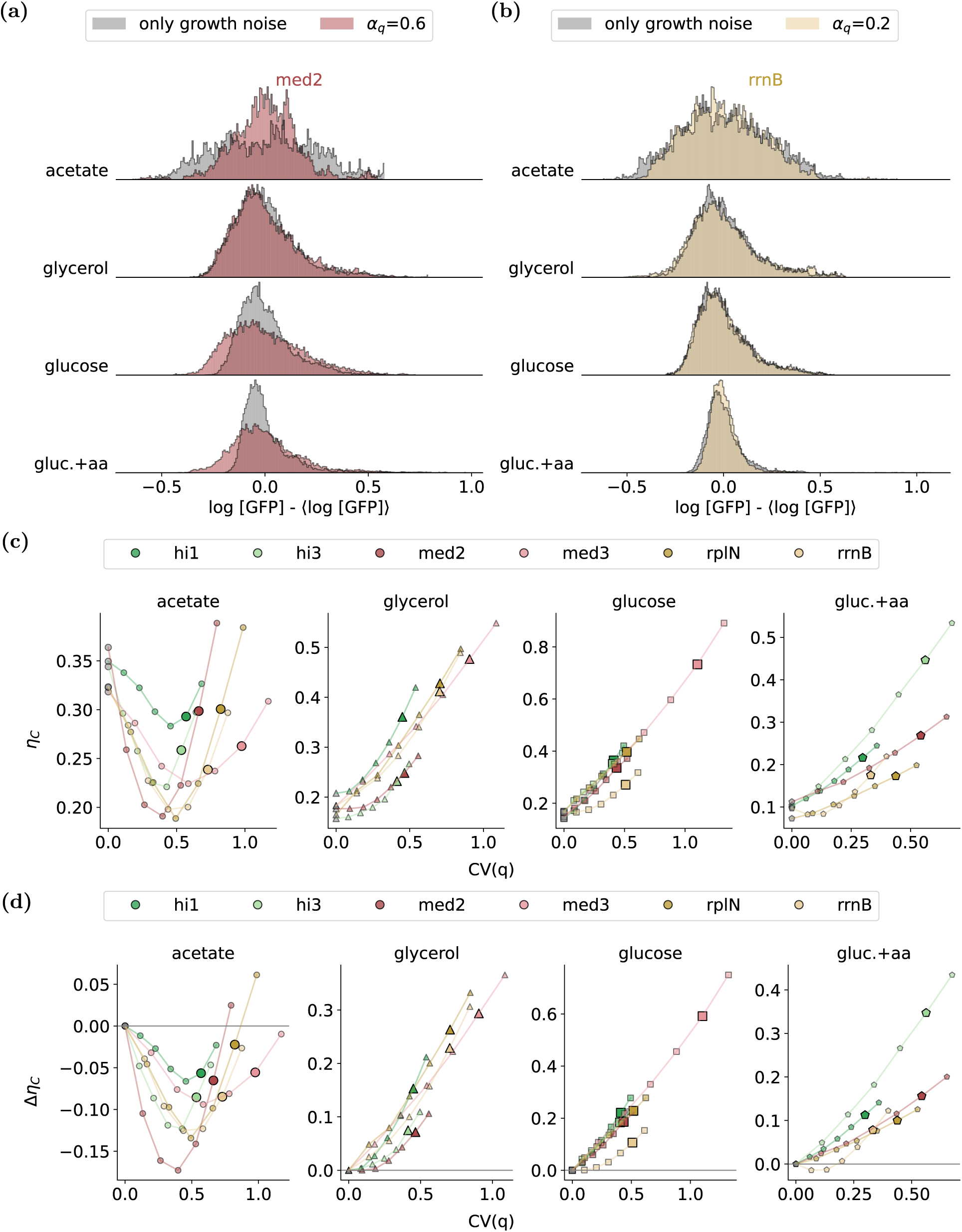
Import of the production noise level. *CV* (*q*) **on concentration fluctuations.** (a) Example distributions for the promoter *med3*. The grey distributions are only driven by growth rate noise, while the red distributions include growth rate noise and production noise, which was scaled by *µ_q_* = 0.6. Thus, in the latter case, the production noise amplitude is 60 % of the experimentally measured noise. In acetate, where growth and production are systematically correlated, scaling the production noise by a factor *α_q_* = 0.6 results in low concentration noise compared to the case where the production noise is absent. (b) For the promoter *rrnB*. In glucose+aa, the production rate of the *rrnB* promoter correlates with the growth rate, similar to the acetate case. The distribution for down-scaled (*α_q_*= 0.2) production noise in glucose+aa is slightly narrower than the distribution without any production noise. (c) The GFP concentration noise level as a function of the production noise level. The markers correspond to values for *α_q_* that range from 0 to 1.2. The large markers correspond to the measured noise levels. (d) Data in (c) replotted, but the noise level of the production noise-free case has been subtracted.

## D Promoter sequences

The following sequences are the promoter sequences as per Sanger sequencing. The GFP-mut2 atg start codon is in smaller letters.

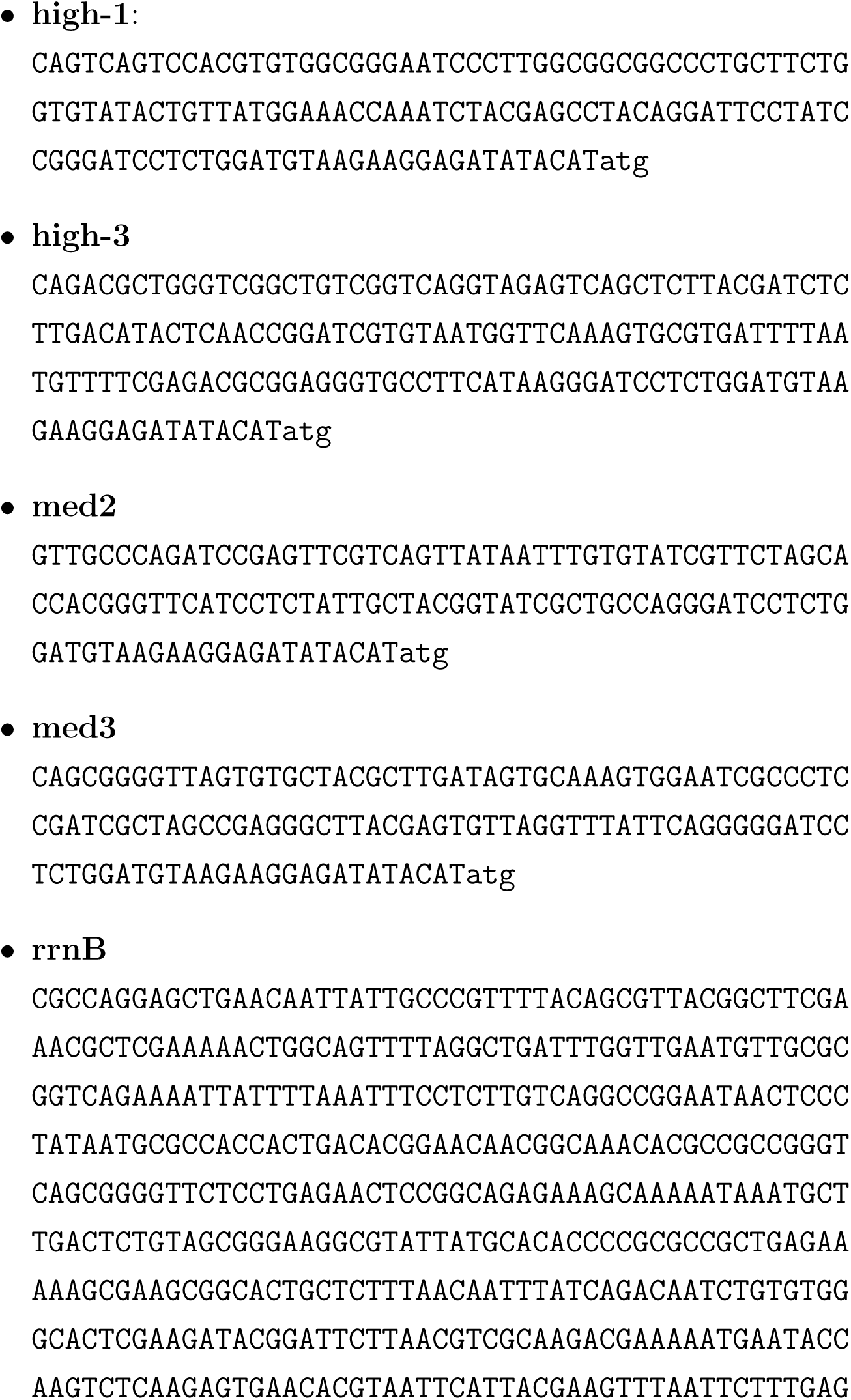
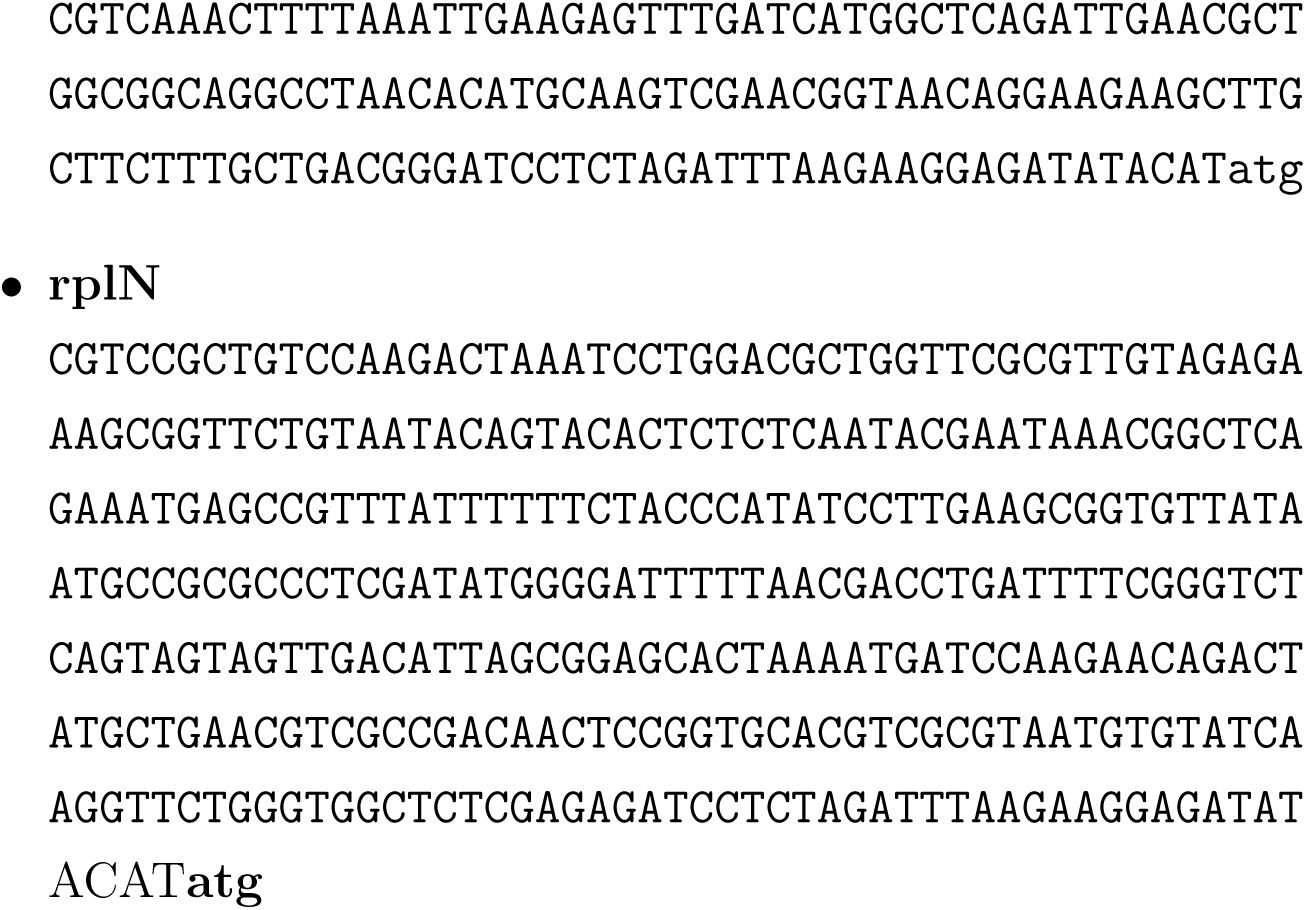

## Notes

https://zenodo.org/records/18129208

### Competing Interest Statement

The authors have declared no competing interest.

